# Cell-autonomous GP130 activation suppresses prostate cancer development via STAT3/ARF/p53-driven senescence and confers an immune-active tumor microenvironment

**DOI:** 10.1101/2024.02.11.579838

**Authors:** Christina Sternberg, Tanja Limberger, Martin Raigel, Karolína Trachtová, Michaela Schlederer, Desiree Lindner, Petra Kodajova, Jiaye Yang, Roman Ziegler, Heidi A. Neubauer, Saptaswa Dey, Torben Redmer, Stefan Stoiber, Václav Hejret, Boris Tichy, Martina Tomberger, Nora S. Harbusch, Simone Tangermann, Monika Oberhuber, Vojtech Bystry, Jenny L. Persson, Sarka Pospisilova, Peter Wolf, Felix Sternberg, Sandra Högler, Sabine Lagger, Stefan Rose-John, Lukas Kenner

**Author notes:** Department of Cell Biology, Charles University, Prague, Czech Republic and Biotechnology and Biomedicine Centre of the Academy of Sciences and Charles University (BIOCEV), Vestec u Prahy, Czech Republic (Roman Ziegler). Institute of Medical Biochemistry, University of Veterinary Medicine Vienna, Vienna, Austria (Heidi A. Neubauer). Contributed equally (Stefan Rose-John, Lukas Kenner).

## Abstract

Prostate cancer ranks as the second most frequently diagnosed cancer in men worldwide. Recent research highlights the crucial roles GP130-mediated signaling pathways play in the development and progression of various cancers, particularly through hyperactivated STAT3 signaling. Here, we find that genetic cell-autonomous activation of the GP130 receptor in prostate epithelial cells triggers active STAT3 signaling and significantly reduces tumor growth *in vivo*. Mechanistically, genetic activation of GP130 signaling mediates senescence via the STAT3/ARF/p53 axis and anti-tumor immunity via recruitment of cytotoxic T-cells, ultimately impeding tumor progression. In prostate cancer patients, high *GP130* mRNA expression levels correlate with better recurrence-free survival, increased senescence signals and a transition from an immune-cold to an immune-hot tumor. Our findings reveal a context-dependent role of GP130/STAT3 in carcinogenesis and a tumor-suppressive function in prostate cancer development. We challenge the prevailing concept of blocking GP130/STAT3 signaling as functional prostate cancer treatment and instead propose cell-autonomous GP130 activation as a novel therapeutic strategy.

## Introduction

Prostate cancer (PCa) is the second most common cancer type diagnosed in men, as reflected by 1.4 million new cases worldwide and 375,000 related deaths in 2020 alone^1^. Corresponding to the vast PCa heterogeneity regarding clinical and molecular features, a wide range of therapeutic approaches is currently in use. The accuracy of these treatments is often hampered by the lack of reliable biomarkers allowing to distinguish aggressive from non-aggressive tumors^2,3^. In search of such biomarkers, aberrant activity of the Glycoprotein 130 kDa (GP130) signaling axis has been identified as a crucial factor in inflammation and carcinogenesis^4,5^. A key downstream mediator of GP130 signaling is the transcription factor Signal transducer and activator of transcription 3 (STAT3)^5^. STAT3 signaling is aberrant in approximately 50% of PCa^6^ and plays a tumor microenvironment (TME)-dependent role in cell proliferation, cell survival, angiogenesis and immune evasion^7–9^. Therefore, a further characterization of the axis connecting GP130 and STAT3 or other potential downstream targets in PCa is important for improved treatment approaches. Other targets activated by GP130 include the Src homology 2 domain-containing tyrosine phosphatase-2 (SHP2), Phosphatidylinositol 3-kinase (PI3K) and the Hippo/YES-associated protein (YAP) pathway^10^, which have themselves been linked with PCa^11–13^. Similarly, the tumor suppressor Phosphatase and tensin homolog (PTEN) is frequently mutated or deleted in PCa^14,15^, thereby eliciting aberrant PI3K activation, contributing to prostate carcinogenesis^16^ and inducing p53-dependent cellular senescence^17,18^. Senescence is a state of cell cycle arrest mediated by the p19^ARF^/p53 or p16^INK4A^/RB pathway and has been shown to inhibit PCa progression^19^. Senescence is often accompanied by the release of inflammatory cytokines, chemokines, growth factors and proteases, referred to as the senescence-associated secretory phenotype (SASP)^20^. The SASP is a double-edged sword exerting tumor-suppressive and tumor-promoting effects. The factors modulating the balance between pro-tumorigenic and anti-tumorigenic senescence effects are likely cell type-specific and not fully understood^21^.

To shed new light on the role of GP130 signaling in PCa pathogenesis and to gain further insight into the complex downstream signaling network, we created a mouse model featuring constitutive, cell-autonomous, and prostate epithelium-specific activation of GP130. Utilizing this model, we aimed at identifying molecular players induced by GP130 signaling and at determining its importance in PCa initiation and progression. Considering recent endeavors to render immune-cold PCa amenable to anti-tumor immunity^22,23^, our study also aimed to elucidate the role of GP130 signaling in shaping the TME, thereby providing valuable insights into its potential use for therapeutic and diagnostic strategies for PCa management.

Our data show that constitutively active GP130 signaling in *Pten*-deficient PCa mice is associated with significantly smaller tumors compared with *Pten*-deficient mice, related with GP130-induced activation of the STAT3/p19^ARF^/p53 tumor suppressor axis mediating senescence. This is accompanied by increased infiltration of cytotoxic T-cells, neutrophils, and macrophages, indicating better anti-tumor defense. These findings are supported by improved survival observed in PCa patients showing high *GP130* mRNA expression, active senescence patterns, and a T-cell mediated anti-tumor immune defense. Together, these results highlight a context-dependent, tumor-suppressive role of GP130/STAT3 signaling in prostate carcinogenesis and suggest that cell-autonomous GP130 activation may be a promising novel therapeutic approach for the treatment of PCa.

## Results

### Constitutive activation of L-gp130 in prostate epithelial cells

To investigate constitutively activated GP130 signaling, we used the Leucine-gp130 (L-gp130) construct introduced by Stuhlman-Laeisz *et al.*^24^ (**Fig. 1a, right panel**), where the entire extracellular part of the wild type GP130 receptor (**Fig. 1a, left panel**) is replaced by a leucine zipper, causing forced receptor dimerization and ligand-independent constitutive activation of downstream signaling. The downstream signaling cascades of L-gp130 include JAK/STAT, PI3K/AKT, MEK/ERK, and Hippo/YAP (**Fig. 1a, right panel**). To study the role of constitutively activated GP130 signaling *in vivo* and to decipher the respective downstream signaling axis involved in PCa, the *L-gp130* transgene composed of a CAG promotor mediating expression of *L-gp130* and *Zoanthus sp.* green fluorescent protein (ZSGreen) was integrated into the ROSA26 locus, which can be transcriptionally activated by Cre-mediated removal of the Westphal stop sequence^4^. We introduced this construct into a conditional mouse model with prostate epithelium-specific Cre expression using Probasin (PB)-Cre4 mice^25,26^. This approach generated mice with a prostate epithelium-specific constitutively active L-gp130 allele (*L-gp130*^peKI/KI^; pe: prostate epithelium). These *L-gp130*^peKI/KI^ mice were then crossed with mice in which *Pten* deletion occurs in the prostate epithelium with sexual maturity, leading to the development of PCa (*Pten*^peΔ/Δ^)^27^. The crossbreed resulted in PCa mice with additional prostate epithelium-specific constitutively activated GP130 signaling (*Pten*^peΔ/Δ^;*L-gp130*^peKI/KI^) (**Fig. 1b**). Deletion of *Pten* and insertion of *L-gp130* were confirmed after the onset of puberty by Polymerase Chain Reaction (PCR) (**Supplementary Fig. 1a**). In addition, prostate epithelium-specific deletion of *Pten* was assessed by immunohistochemistry (IHC) analysis of phospho-AKT (p-AKT) levels, as loss of PTEN leads to phosphorylation of AKT^16^. *Pten*^peΔ/Δ^ and *Pten*^peΔ/Δ^;*L-gp130*^peKI/KI^ showed elevated p-AKT expression compared to wild type and *L-gp130*^peKI/KI^ prostate samples (**Fig. 1c, upper panel**). Further confirmation of prostate-specific deletion of *Pten* was obtained by Western blot analysis and quantification of p-AKT levels. *Pten*^peΔ/Δ^ and *Pten*^peΔ/Δ^;*L-gp130*^peKI/KI^ showed significantly increased p-AKT expression compared to wild type and *L-gp130*^peKI/KI^ prostate samples, whereas total-AKT (t-AKT) expression was not affected between the different genotypes (**Supplementary Fig. 1b-c**). To confirm the functional expression of the *L-gp130* construct, we examined ZSGreen expression via immunofluorescence (IF). We found ZSGreen expression in *L-gp130*^peKI/KI^ and *Pten*^peΔ/Δ^;*L-gp130*^peKI/KI^ but not in wild type and *Pten*^peΔ/Δ^ prostates (**Fig. 1c**). Of note, endogenous wild type *Gp130* mRNA levels were not changed by *L-gp130* expression in the prostate (**Supplementary Fig. 1d**). Overall, we generated a mouse model allowing us to examine the consequences of cell-autonomous, prostate epithelial cell-specific GP130 signaling.

**Fig. 1:**
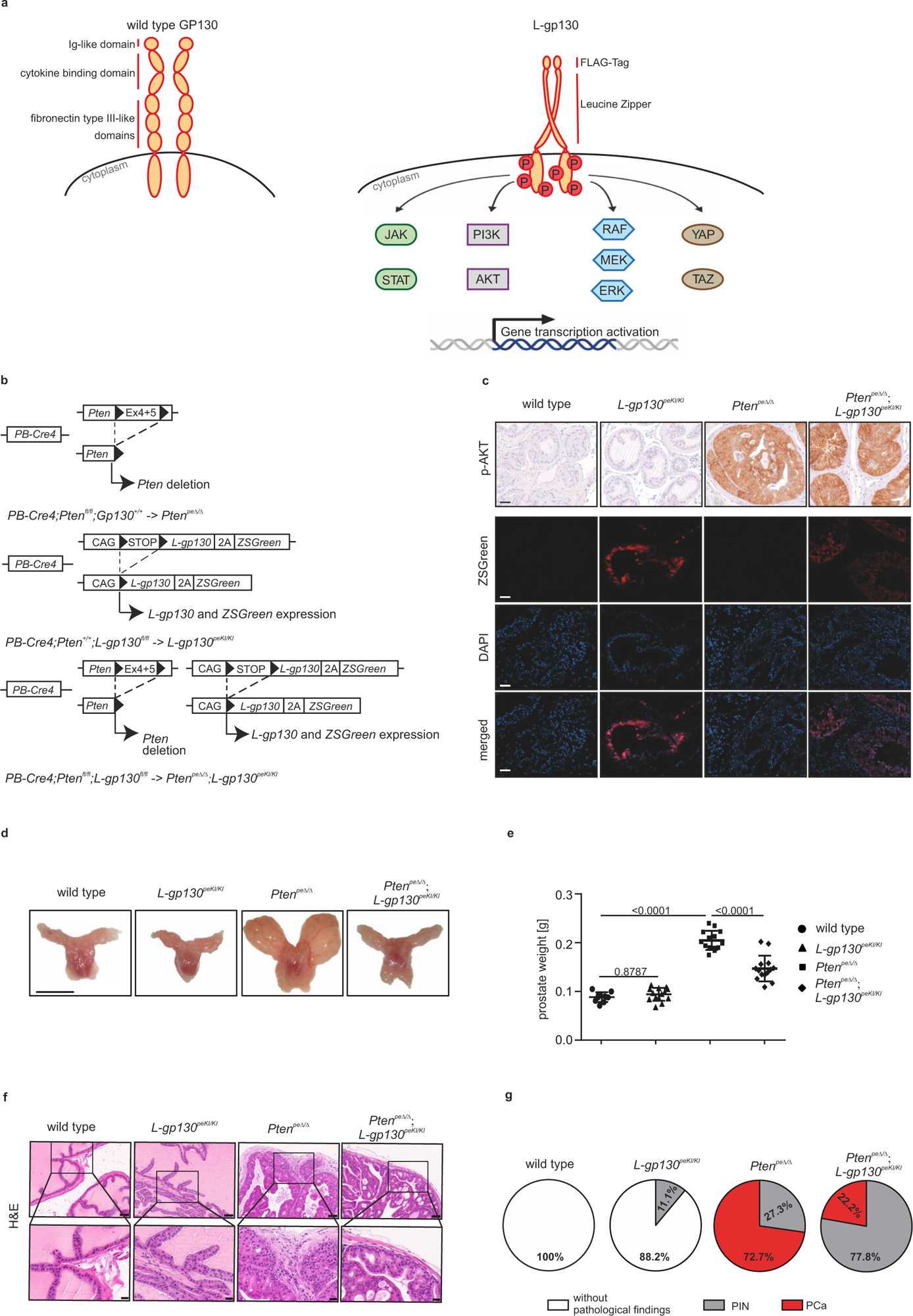
Prostate epithelium-specific genetic deletion of *Pten* and insertion of *L-gp130* reduces progressive prostate tumorigenesis. a) Illustration of wild type GP130 receptor (left) and Leucine-gp130 (L-gp130) construct (right). The wild type GP130 receptor consists of an extracellular domain comprising an Ig-like domain, a cytokine binding domain, three fibronectin type III-like domains, a transmembrane domain, and a cytoplasmic domain. For generating L-gp130, wild type GP130 was truncated 15 amino acids above the transmembrane domain and replaced by the leucine zipper region of the human c-JUN gene and a FLAG-Tag. L-gp130 expression can activate several downstream signaling cascades: JAK/STAT, RAF/MEK/ERK, PI3K/AKT and Hippo/YAP signaling. P: phosphorylation b) Illustration of the genetic approach for conditional deletion of *Pten* (exon 4+5) or/and insertion of *L-gp130-ZSGreen* in prostate epithelial cells under the control of the androgen-regulated and prostate-specific Probasin (PB) promoter after *Cre-*mediated recombination resulting in PB-Cre4;*Pten*^fl/fl^;*L-gp130*^+/+^ mice (hereafter *Pten*^peΔ/Δ^), PB-Cre4;*Pten*^+/+^;*L-gp130*^fl/fl^ mice (hereafter *L-gp130*^peKI/KI^) and PB-Cre4;*Pten*^fl/fl^;*L-gp130*^fl/fl^ mice (hereafter *Pten*^epΔ/Δ^;*L-gp130*^peKI/KI^). pe: prostate epithelium; fl: floxed site; ex: exon; 2A: 2A peptide; CAG: CAG promoter; KI: knock in; Δ: knock out; c) Representative immunohistochemistry (IHC) pictures of phospho-AKT (p-AKT) and immunofluorescence (IF) pictures of co-stainings of ZSGreen (red) and DAPI (blue) in prostates from wild type, *L-gp130*^peKI/KI^, *Pten*^peΔ/Δ^, and *Pten*^peΔ/Δ^;*L-gp130*^peKI/KI^ mice. DAPI is used as a nuclear stain. Scale bar: 40 µm. d) Gross anatomy of representative prostates isolated from wild type, *L-gp130*^peKI/KI^, *Pten*^peΔ/Δ^, and *Pten*^peΔ/Δ^;*L-gp130*^peKI/KI^ mice. Scale bar: 1 cm. e) Prostate weight of wild type (n=10), *L-gp130*^peKI/KI^ (n=14), *Pten*^peΔ/Δ^ (n=14), and *Pten*^peΔ/Δ^;*L-gp130*^peKI/KI^ (n=14) mice. Individual biological replicates are shown. Data are plotted as the means±SD and p-values were determined by ordinary one-way ANOVA with Tukey’s multiple comparisons test. f) Representative pictures of H&E stains of mouse prostates from the indicated genotypes at low (top) and high (bottom) magnification. Scale bar upper panel: 60 µm, scale bar lower panel: 20 µm. g) Quantification of histopathological analysis of prostate tissue from wild type (n=9), *L-gp130*^peKI/KI^ (n=9), *Pten*^peΔ/Δ^ (n=11), and *Pten*^peΔ/Δ^;*L-gp130*^peKI/KI^ (n=9) mice in regards of histomorphological criteria for aggressive growth patterns: without pathological findings (white); PIN: prostate intraepithelial neoplasia (grey); PCa: prostate cancer (red).

### Constitutively active GP130 signaling reduces *Pten*-deficient tumor growth *in vivo*

We next examined the impact of constitutively active GP130 signaling in 19-week old mice and found that prostates of wild type and *L-gp130*^peKI/KI^ mice were macroscopically indistinguishable. As expected, *Pten*^peΔ/Δ^ mice developed grossly visible PCa (**Fig. 1d**). Intriguingly, mice with concomitant activation of GP130 signaling (*Pten*^peΔ/Δ^;*L-gp130*^peKI/KI^) developed smaller prostate tumors compared to *Pten*-deficient mice, resulting in a significantly reduced prostate weight of *Pten*^peΔ/Δ^;*L-gp130*^peKI/KI^ compared to *Pten*^peΔ/Δ^ mice (**Fig. 1d-e**). Assessment of hematoxylin and eosin (H&E)-stained murine prostates (**Fig. 1f**) revealed no pathological features in wild type and *L-gp130*^peKI/KI^ mice, except for one *L-gp130*^peKI/KI^ animal (accounting for 11.1% of analyzed mice) (**Fig. 1g**). This animal exhibited prostatic intraepithelial neoplasia (PIN), a precursor for PCa. The vast majority (72.7%) of the *Pten*^peΔ/Δ^ mice showed PCa^28^, whereas only 22.2% of *Pten*^peΔ/Δ^;*L-gp130*^peKI/KI^ mice developed PCa. Instead, 77.8% exhibited only PIN, displaying a less aggressive morphology compared to *Pten*^peΔ/Δ^ mice. These findings support a tumor-suppressive role of GP130 signaling in PCa *in vivo*.

### *L-gp130* expression in prostate epithelial cells enriches STAT3 target gene expression

To unravel molecular gene expression patterns associated with the observed phenotypes and to elucidate which of the possible downstream signaling axes is activated upon *L-gp130* insertion, we performed RNA sequencing (RNA-Seq) analysis of prostate tissue from wild type, *L-gp130*^peKI/KI^, *Pten*^peΔ/Δ^ and *Pten*^peΔ/Δ^;*L-gp130*^peKI/KI^ mice. As our mouse model allows prostate epithelium-specific modulation and to specifically isolate these prostate epithelial cells, we sorted cells obtained from prostate tissue by magnetic bead-based cell sorting for EpCAM, a marker for epithelial cells that is expressed uniformly across all four genotypes (**Fig. 2a**). This prostate epithelial fraction was then subjected to RNA-Seq as previously described^29^ (**Fig. 2b**). Clustering of the samples by 3D-principal component analysis based on gene expression revealed that individual replicates clustered within the genotypes and confirmed different transcription profiles (**Supplementary Fig. 2a**). Differential gene expression analysis of *Pten*^peΔ/Δ^ and *Pten*^peΔ/Δ^;*L-gp130*^peKI/KI^ prostate epithelial cells showed significant upregulation of 807 and downregulation of 475 genes in *Pten*^peΔ/Δ^;*L-gp130*^peKI/KI^ prostates (**Fig. 2c**). Notably, we detected nearly twice as many upregulated genes as downregulated genes, which supports the idea that constitutively active GP130 serves as a central receptor of signal transduction and activator of transcription. There was also a considerable, albeit smaller number of 470 genes that were significantly upregulated when comparing *L-gp130*^peKI/KI^ and wild type prostate epithelial cells, as well as 335 downregulated genes (**Supplementary Fig. 2b**). Fast pre-ranked gene set enrichment analysis (fGSEA) of the “Prostate cancer” gene set from the Kyoto Encyclopedia of Genes and Genomes (KEGG) collection from the Molecular signature database (MSigDB)^30,31^ showed a significant downregulation of analyzed genes in *Pten*^peΔ/Δ^;*L-gp130*^peKI/KI^ compared to *Pten*^peΔ/Δ^ samples (**Fig. 2d**), which supports our mouse data showing smaller tumors in *Pten*^peΔ/Δ^;*L-gp130*^peKI/KI^ mice.

**Fig. 2:**
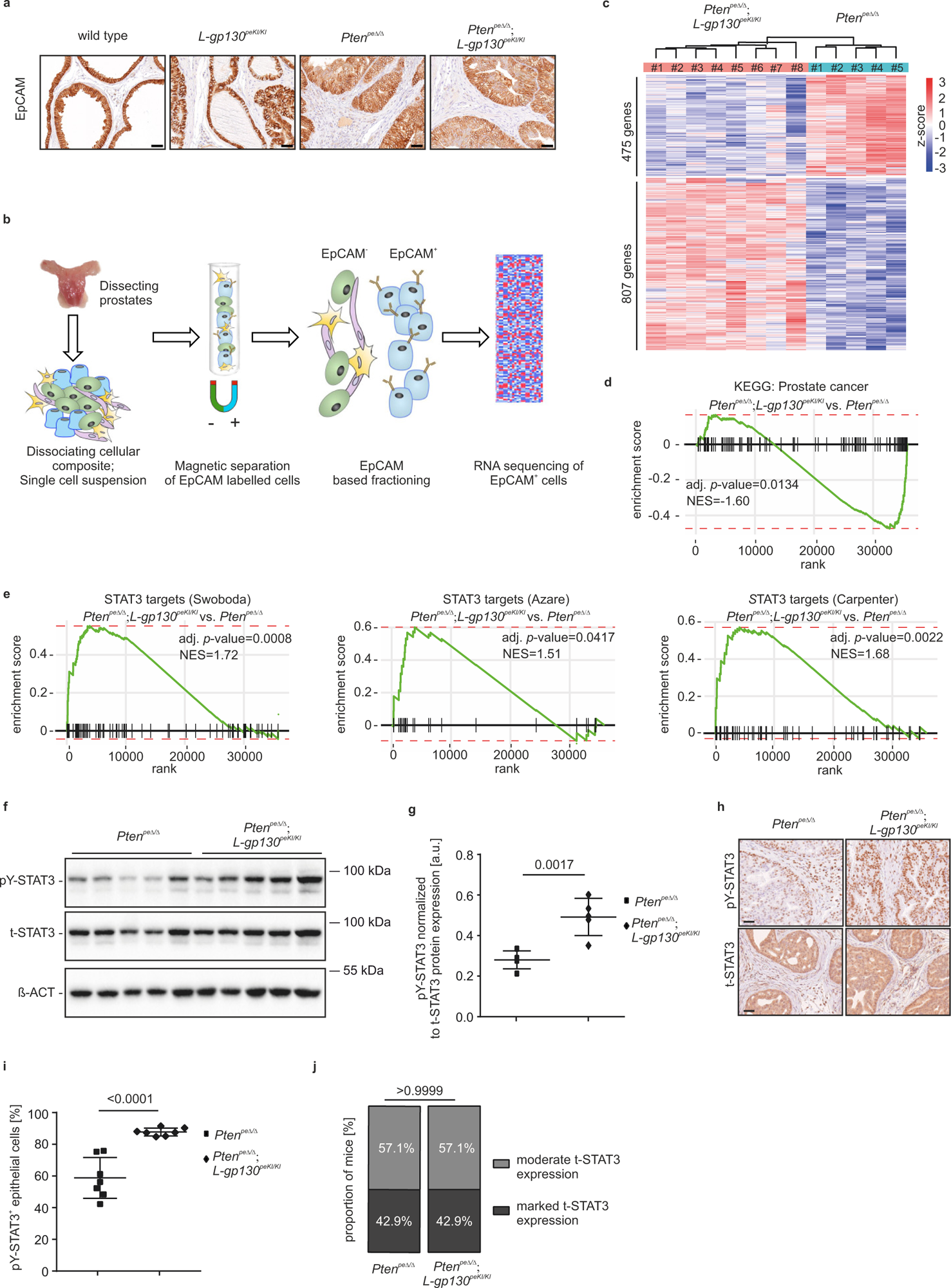
Prostate-specific, cell-autonomous insertion of *L-gp130* leads to activation of the STAT3 transcription factor and STAT3 target gene expression. a) Representative IHC pictures of mouse prostates from wild type, *L-gp130*^peKI/KI^, *Pten*^peΔ/Δ^ and *Pten*^peΔ/Δ^;*L-gp130*^peKI/KI^ mice stained for the epithelial marker EpCAM. Scale bar: 40 µm b) RNA-Seq workflow showing processing and magnetic bead-based enrichment of EpCAM-positive (EpCAM^+^) mouse prostate tissue. Prostates were dissected and enzymatically and mechanically dissociated to generate single cell suspensions. Cells were labelled with biotinylated anti-EpCAM antibody and enriched from the bulk population using streptavidin-coated magnetic beads. EpCAM^+^ cells were subjected to RNA-Seq analysis. c) Heatmap and number of differentially expressed genes (log2norm) based on adj. p-value ≤0.05 and fold change ≥2 cut-off values comparing *Pten*^peΔ/Δ^ and *Pten*^peΔ/Δ^;*L-gp130*^peKI/KI^ prostate epithelial cells (n≥5). blue: downregulated, red: upregulated. d) Fast pre-ranked gene set enrichment analysis (fGSEA) of the KEGG gene set “Prostate cancer” with genes regulated in *Pten*^peΔ/Δ^;*L-gp130*^peKI/KI^ compared to *Pten*^peΔ/Δ^ prostate epithelial cells. Genes sorted based on their Wald statistics are represented as vertical lines on the x-axis. NES: normalized enrichment score. e) Fast pre-ranked gene set enrichment analysis (fGSEA) of three previously published STAT3 target signatures (“STAT3 targets (Swoboda)”, “STAT3 targets (Azare)”, “STAT3 targets (Carpenter)”) with genes regulated in *Pten*^peΔ/Δ^;*L-gp130*^peKI/KI^ compared to *Pten*^peΔ/Δ^ prostate epithelial cells. Genes sorted based on their Wald statistics are represented as vertical lines on the x-axis. NES: normalized enrichment score. f) Western Blot analysis of prostate protein lysates for phosphoTyrosine705-STAT3 (pY-STAT3) and total-STAT3 (t-STAT3) expression in *Pten*^peΔ/Δ^ and *Pten*^peΔ/Δ^;*L-gp130*^peKI/KI^ mice (n=5). β-ACTIN (β-ACT) served as loading control. g) Quantification of pY-STAT3 protein levels relative to t-STAT3 protein levels shown in f). h) Representative pictures of immunohistochemistry (IHC) staining of pY-STAT3 and t-STAT3 expression in prostate sections of *Pten*^peΔ/Δ^ and *Pten*^peΔ/Δ^;*L-gp130*^peKI/KI^ mice. Scale bar: 40 µm. i-j) Quantitative analysis of pY-STAT3 (i) and semi-quantitative analysis of t-STAT3 (j) IHC stainings shown in g) (n=7). (g,i-j) Individual biological replicates are shown (g,i). Data are plotted as the means±SD and p-values were determined by unpaired two-tailed Student’s t-tests (g,i) or Mann-Whitney test (j).

To determine which downstream signaling cascade is activated by L-gp130 in PCa, we performed fGSEA of *Pten*^peΔ/Δ^;*L-gp130*^peKI/KI^ compared to *Pten*^peΔ/Δ^ samples using the REACTOME gene set collection derived from MSigDB. We detected no apparent change in the regulation of PI3K/AKT, RAS/RAF/ERK/MAPK or Hippo/YAP signaling cascades (**Supplementary Fig. 2c**). Interestingly, we found significantly upregulated STAT3 target genes by analyzing three independent, previously described sets of STAT3 target genes^32–34^. These results imply that *L-gp130* expression correlated with STAT3 activity, which in turn acts as a transcription factor in PCa (**Fig. 2e**). In accordance, Western blot analysis also showed activation of STAT3, reflected in increased phosphoY705-STAT3 (pY-STAT3) levels and unaltered total-STAT3 (t-STAT3) levels in *Pten*^peΔ/Δ^;*L-gp130*^peKI/K^ compared to *Pten*^peΔ/Δ^ prostates (**Fig. 2f-g**). This finding was further confirmed using IHC, which showed a nearly 50% increase in pY-Stat3 positive cells in *Pten*^peΔ/Δ^;*L-gp130*^peKI/KI^ compared to *Pten*^peΔ/Δ^ samples, while t-STAT3 levels assessed semi-quantitatively were constant in both genotypes (**Fig. 2h-j**). This increase is on top of the already nearly 60% pY-STAT3^+^ epithelial cells seen in *Pten*^peΔ/Δ^ samples. We additionally observed an upregulation of STAT3 target genes (**Supplementary Fig. 2d**) and of pY-STAT3 abundance (**Supplementary Fig. 2e-i**) when comparing *L-gp130*^peKI/KI^ and wild type prostate epithelial cells. Of note, in the stromal cells of mice with *L-gp130* insertion in the prostate epithelium (*L-gp130^peKI/KI^* and *Pten^peΔ/Δ^;L-gp130^peKI/KI^* mice), we detected no increase in pY-STAT3 levels compared to wild type and *Pten*^peΔ/Δ^ mice, respectively (**Supplementary Fig. 2j**). This underscores the prostate-epithelium specificity of our mouse model. Taken together, L-gp130 expression in prostate epithelial cells activates STAT3 signaling, as evidenced by significantly upregulated STAT3 target genes and increased pY-STAT3 levels.

### GP130 expression correlates with prolonged survival and STAT3 expression in PCa patients

Based on our mouse data, we hypothesized that high GP130 expression in PCa patients would correlate with favorable clinical outcomes. To investigate the relationship between *GP130* mRNA expression levels and PCa, we examined the TCGA-PRAD cohort^35^ comprising primary PCa patients and found a significant decrease in *GP130* mRNA expression in prostate tumor tissue compared to adjacent healthy tissue (**Fig. 3a**). Stratifying patients based on their *GP130* mRNA demonstrated that high *GP130* expression is linked to a greater probability of disease-free survival compared to low *GP130* expression (**Fig. 3b**). In the TCGA-PRAD cohort, 7.2% of patients with prostate adenocarcinoma have alterations in the *GP130* gene, with 6.3% accounting for deep deletions and 0.9% for missense mutations of unknown significance (**Fig. 3c**). We therefore hypothesized that mutations leading to altered *GP130* expression could impact the initiation and/or progression of PCa. To test the validity of our findings, we analyzed four additional data sets from the Oncomine platform^36^. The results support a significant decrease in *GP130* mRNA expression in PCa compared to normal prostate glands, highlighting the potential impact of *GP130* alterations on PCa (**Fig. 3d**). We also observed a significant reduction in *GP130* expression relative to the primary PCa site during PCa progression in recurrent and advanced PCa and metastasis (**Supplementary Fig. 3a**). Using the SurvExpress Analysis webtool^37^, we next examined the MSKCC Prostate GSE21032 data set by Taylor *et al.*^38^ in terms of survival, as it provides not only data on primary but also metastatic and recurrent PCa. We assessed risk groups by a median split of samples based on their prognostic index and observed high expression of *GP130* in low-risk PCa patients and *vice versa* (**Supplementary Fig. 3b**). In support of our previous findings from the TCGA-PRAD cohort, correlating biochemical recurrence-free survival time with *GP130* mRNA expression levels, we detected a significantly higher probability of biochemical recurrence-free survival associated with high as compared to low *GP130* levels (**Supplementary Fig. 3c**). These findings corroborate a tumor suppressive role of *GP130* expression in PCa. Interestingly, *GP130* mRNA expression positively correlated with *STAT3* expression in PCa patients in both the TCGA-PRAD cohort and Taylor data set (MSKCC Prostate GSE21032), providing further evidence of the interconnected signaling between GP130 and STAT3 in PCa (**Fig. 3e** and **Supplementary Fig. 3d**). Together, these human patient data suggest that *GP130* expression could serve as a useful marker to stratify PCa cases into low- and high-risk groups.

**Fig. 3:**
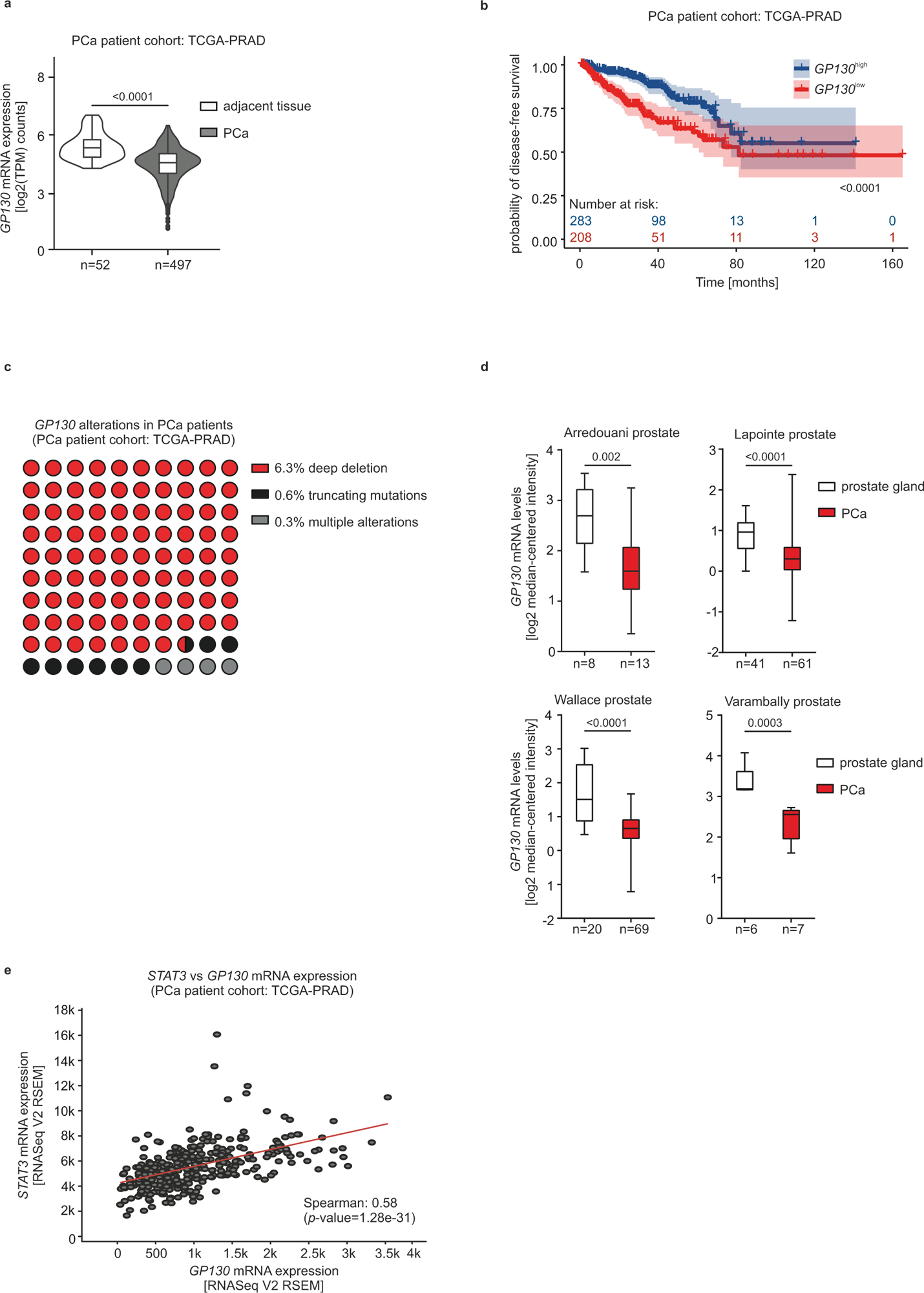
High *GP130* gene expression is significantly associated with low-risk human PCa groups and with better recurrence-free survival in human PCa. a) *GP130* gene expression in adjacent (n=52) and PCa (n=497) tissue in TCGA-PRAD data set. Statistical analysis of the two risk groups was determined by using the Mann-Whitney test. b) Kaplan-Meier plot showing time of disease-free survival in months for *GP130*^low^ and *GP130*^high^ risk groups of the TCGA-PRAD data set. Groups were assessed based on the maximally selected rank statistics. blue: high *GP130* expressing group, red: low *GP130* expressing group. The blue and red numbers below horizontal axis represent the number of patients. c) Proportion of *GP130* alterations in the TCGA-PRAD data set. Mutation types: deep deletion (n=21; red), truncating mutations (n=2; black) and multiple alterations (n=1; grey). One patient has simultaneous mutations. The data originate from cBioPortal. d) *GP130* mRNA expression levels of four different data sets of PCa patient samples compared to healthy prostate sample control. Normalized data and statistical analyses were extracted from the Oncomine Platform. The respective prostate data set and n-numbers are indicated. Representation: boxes as interquartile range, horizontal line as the mean, whiskers as lower and upper limits. e) Spearman-correlation analysis of *GP130* and *STAT3* expression in TCGA-PRAD data set using cBioPortal analysis tool.

### L-gp130 promotes STAT3/p19^ARF^/p53-induced senescence upon *Pten*-loss

To further understand the molecular mechanisms underlying the observed reduction in tumor size in mice expressing constitutively active *Gp130* in the prostate epithelium, we investigated changes in gene expression. Performing fGSEA using the HALLMARK gene set collection from MSigDB, we identified significantly deregulated gene sets that rely on L-gp130 expression in prostate tumorigenesis. Upon *L-gp130* insertion, the “IL-6/JAK/STAT3 signaling” gene set was upregulated in *L-gp130*^peKI/KI^ and *Pten*^peΔ/Δ^;*L-gp130*^peKI/KI^ mice, compared to wild type and *Pten*^peΔ/Δ^ mice, respectively (**Fig. 4a** and **Supplementary Fig. 4a**). This is noteworthy as the cytokine Interleukin-6 (IL-6) activates the Janus kinase (JAK) and subsequently STAT3 by binding to the GP130 receptor^10^, and therefore upregulated ‘IL-6/JAK/STAT3 signaling’ aligns with our previous results on the induction of the STAT3 signaling cascade. From all HALLMARK gene sets, we found 32 being significantly deregulated when comparing *Pten*^peΔ/Δ^;*L-gp130*^peKI/KI^ and *Pten*^peΔ/Δ^ samples. Among these gene sets, the downregulated HALLMARK gene set “Androgen response” (**Fig. 4a**) points to less androgen receptor signaling in *Pten*^peΔ/Δ^;*L-gp130*^peKI/KI^ mice, which is in line with its crucial role in PCa development^39^ and the observed reduction in cancer aggressiveness in our *Pten*^peΔ/Δ^;*L-gp130*^peKI/KI^ mice. The two HALLMARK gene sets showing the most pronounced downregulation were „Oxidative phosphorylation” and „Fatty acid metabolism“. This observation is noteworthy as we have previously demonstrated an inverse association between the regulation of oxidative phosphorylation and the TCA cycle with STAT3 expression^40,41^. This downregulation is also seen in the corresponding KEGG and Biological Processes from Gene Ontology pathways (GO-BP) gene sets (**Supplementary Fig. 4b**). Given that the downregulation of these pathways has previously been shown to rely on STAT3 signaling, it underscores the importance of active STAT3 signaling in the context of this study. Surprisingly, the two most prominent upregulated gene sets are the proliferation-associated „MYC targets V1” and „MYC targets V2” (**Fig. 4a**). We also observed an upregulation of MYC target genes in the comparison of *L-gp130*^peKI/KI^ and wild type (**Supplementary Fig. 4a**). As *MYC* gene expression is regulated by GP130/STAT3^42^, this might contribute to the observed upregulation of MYC target genes in both comparisons (*Pten*^peΔ/Δ^;*L-gp130*^peKI/KI^ versus *Pten*^peΔ/Δ^ and *L-gp130*^peKI/KI^ versus wild type). Additionally, the cell cycle-related gene sets „E2F targets” and „G2M checkpoint” were significantly upregulated. Considering the reported potential of the IL-6/STAT3 axis to drive rather than inhibit tumor cell proliferation^43^, we investigated proliferation. We did not observe a significant difference in Ki67 assessed by IHC staining (**Fig. 4b-c**). Another gene set that was observed to be significantly upregulated is the „P53 pathway“, which is known to mediate oncogene-induced senescence in prostate tumorigenesis^44,45^ and in the Pten-deficient PCa context^9^. Therefore we hypothesized that the induction of senescence in *Pten*^peΔ/Δ^;*L-gp130*^peKI/KI^ compared to *Pten*^peΔ/Δ^ mice causes the smaller tumors observed in *Pten*^peΔ/Δ^;*L-gp130*^peKI/KI^ mice. Senescence is often accompanied by the upregulation of promyelocytic leukemia protein (PML)^46^. Indeed, we observed increased numbers of PML nuclear bodies in our *Pten*^peΔ/Δ^;*L-gp130*^peKI/KI^ mice (**Fig. 4d-e**). An additional defining characteristic of senescent cells is the release of inflammatory cytokines and signaling molecules referred to as SASP^20^. To investigate the alteration of SASP-related genes in *Pten*^peΔ/Δ^;*L-gp130*^peKI/KI^ prostate epithelial cells, we performed fGSEA using the “Core SASP of Pten-loss induced cellular senescence (PICS)”^47^ gene set, previously described to be induced upon PICS, and found it to be significantly upregulated in *Pten*^peΔ/Δ^;*L-gp130*^peKI/KI^ mice compared to *Pten*^peΔ/Δ^ mice (**Supplementary Fig. 4c**).

**Fig. 4:**
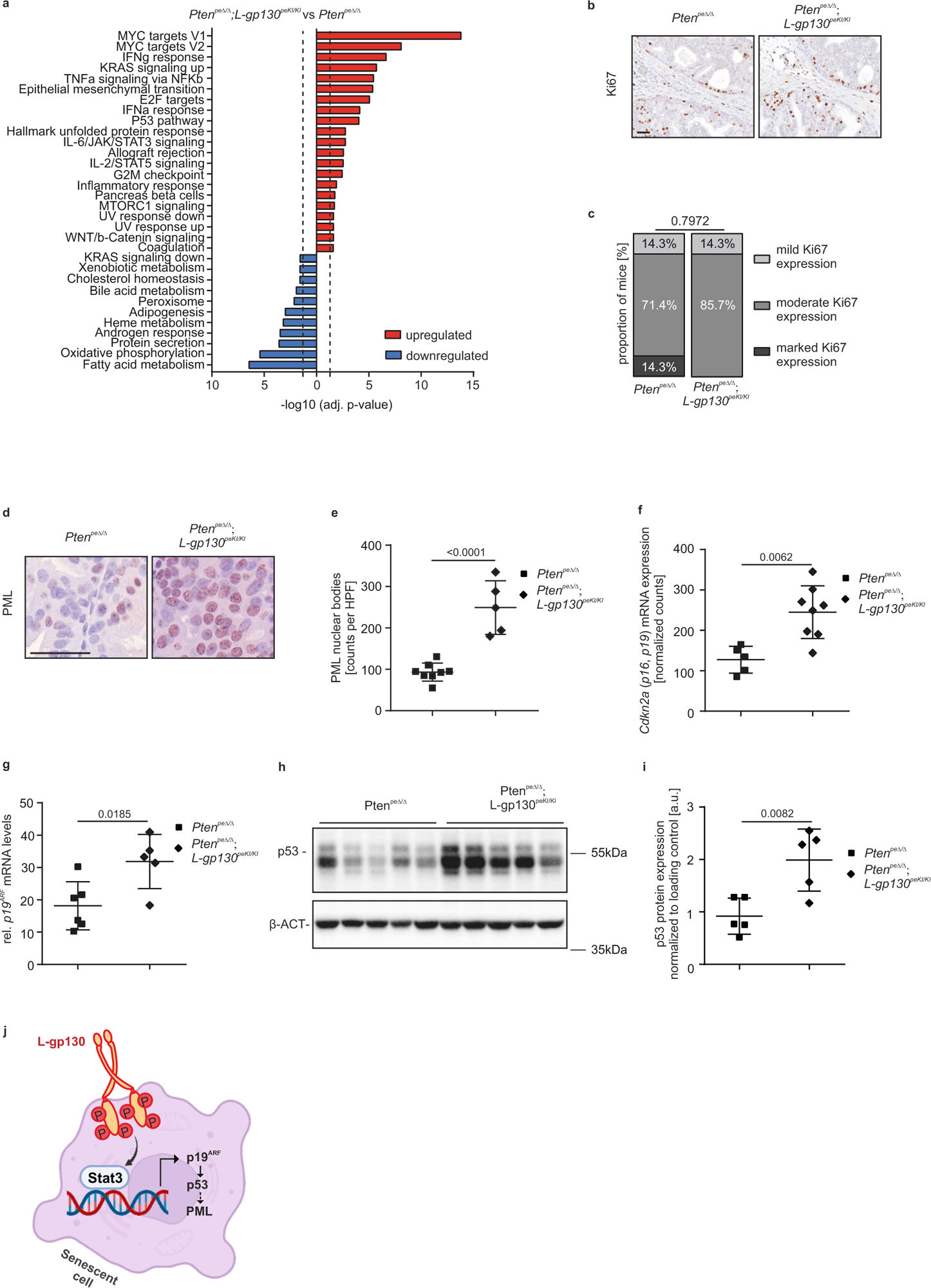
Expression of L-gp130 induces p19^ARF^-p53-driven senescence in *Pten*-deficient PCa. a) Fast pre-ranked gene set enrichment analysis (fGSEA) of significantly enriched HALLMARK gene sets with genes regulated in *Pten*^peΔ/Δ^;*L-gp130*^peKI/KI^ compared to *Pten*^peΔ/Δ^ prostate epithelial cells. Dotted line: adj. p-value (-log10(0.05)), blue: downregulated, red: upregulated; b) Representative pictures of immunohistochemistry (IHC) staining of mouse prostates from the indicated genotypes stained for the proliferation marker Ki67. Scale bar: 40 µm. c) Semi-quantitative analysis of Ki67^+^ prostate epithelial cells in the indicated genotypes (n=7) shown in b). d) Representative pictures of immunohistochemistry (IHC) staining of PML of *Pten*^peΔ/Δ^ and *Pten*^peΔ/Δ^;*L-gp130*^peKI/KI^ prostates. Scale bar: 40 µm. e) Quantification of PML nuclear bodies per high power field (HPF) shown in d) (n≥5). f) *Cdkn2a* mRNA expression levels in *Pten*^peΔ/Δ^ and *Pten*^peΔ/Δ^;*L-gp130*^peKI/KI^ prostates (n≥5). g) qPCR mRNA expression analysis of *p19^ARF^* in mouse prostate tissue of *Pten*^peΔ/Δ^ and *Pten*^peΔ/Δ^;*L-gp130*^peKI/KI^ mice (n≥5). Signals are relative to the geometric mean of housekeeping genes. h) Western Blot analysis of prostate protein lysates of *Pten*^peΔ/Δ^ and *Pten*^peΔ/Δ^;*L-gp130*^peKI/K^ mice (n=5) for p53 expression. β-ACTIN (β-ACT) served as loading control. i) Quantification of p53 protein levels shown in h) normalized to loading control. j) Proposed model of GP130 signaling induced senescence. L-gp130 activated STAT3 binds to its binding sites in Cdkn2a promoter, followed by upregulation of p19^ARF^ and p53 expression promoting senescence in PCa. (c,e-g,i) Individual biological replicates are shown (e-g,i). Data are plotted as the means±SD and p-values were determined by Mann-Whitney test (c,f), unpaired two-tailed Student’s t-tests (e,g,i).

Upon closer examination of the molecular players involved in senescence induction, we found that the p19^ARF^/p53-dependent pathway was activated. We observed significantly enhanced expression of *Cdkn2a* mRNA, which encodes both *p16^INK4A^* and *p19^ARF^* (**Fig. 4f**)^48^. Using *p19^ARF^* specific primers revealed a significant upregulation of this previously described STAT3 target gene^9^ (**Fig. 4g**). We observed a significant increase in p53 protein abundance (**Fig. 4h-i**). In line with this, we also noted that gene sets representing transcriptional p53 activity were significantly upregulated in our fGSEA analysis of GO-BP gene sets derived from MSigDB (**Supplementary Fig. 4d**). Based on our data, we thus propose a model of GP130 signaling-induced senescence in PCa, in which STAT3, activated by L-gp130, upregulates *p19^ARF^* mRNA expression, followed by increased p53 expression and induction of senescence as seen by elevated PML expression (**Fig. 4j**).

### GP130 signaling recruits anti-tumor infiltrating immune cells

Senescence is closely connected to the TME, known to be immune-cold in PCa^22,23^. Consequently, we focused our examination on the impact of constitutively active GP130 on the TME. Indeed, upon analysis of H&E stained prostate sections we detected high-grade immune cell infiltration in 66.7% of *Pten*^peΔ/Δ^;*L-gp130*^peKI/KI^ mice compared to 36.4% of *Pten*^peΔ/Δ^ mice (**Fig. 5a**). Only few immune cells were seen in wild type and *L-gp130*^peKI/KI^ prostates (**Supplementary Fig. 5a**). IHC analysis allowed us to examine the infiltrating immune cell subtypes and their distribution and revealed a significantly higher number of CD45^+^ cells in *Pten*^peΔ/Δ^;*L-gp130*^peKI/KI^ mice compared to *Pten*^peΔ/Δ^ mice and the occurrence of minimal CD45^+^ cells in wild type and *L-gp130*^peKI/KI^ mice (**Fig. 5b-c** and **Supplementary Fig. 5b-c**).

**Fig. 5:**
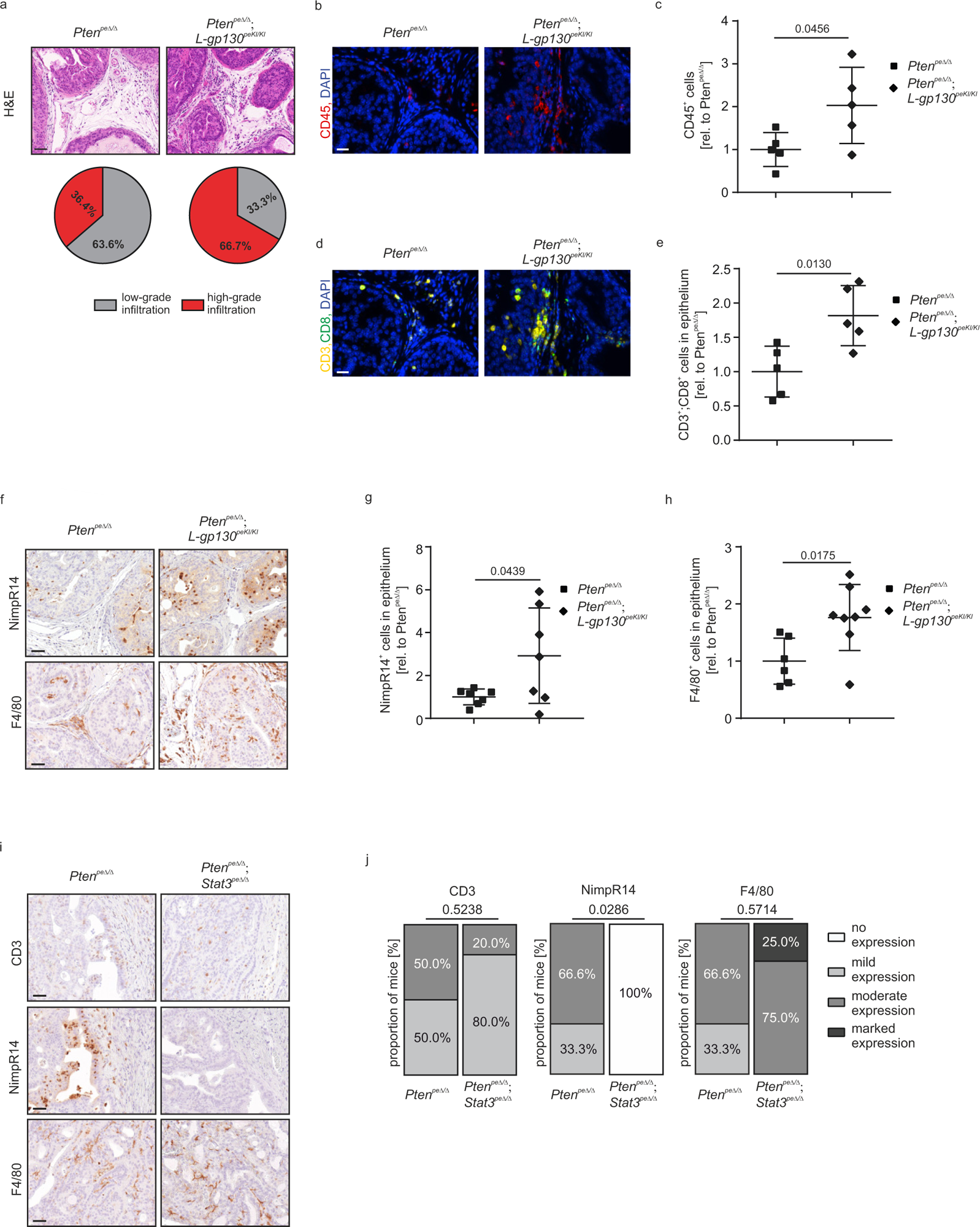
Expression of L-gp130 in *Pten*^peΔ/Δ^ mice increases infiltration of CD8^+^ T-cells mediating anti-tumor defense. a) Representative pictures of H&E stains (upper panel) showing immune infiltrate and quantification of histopathological analysis (lower panel) of prostate tissue from *Pten*^peΔ/Δ^ (n=11) and *Pten*^peΔ/Δ^;*L-gp130*^peKI/KI^ (n=9) mice in regards of infiltration (low-grade (grey) and high-grade (red)). Scale bar: 60 µm. b) Representative pictures of immunofluorescence (IF) staining of CD45 (red) and DAPI (blue) of mouse prostates with indicated genotypes. DAPI is used as a nuclear stain. Scale bar: 20 µm. c) Quantification of CD45^+^ cells of IF stainings shown in b) (n=5). A whole slide scan of stained prostate tissue per mouse was analyzed. The percentage of positive cells relative to *Pten*^peΔ/Δ^ was calculated. d) Representative pictures of immunofluorescence (IF) staining of CD3 (yellow), CD8 (green) and DAPI (blue) of mouse prostates with indicated genotypes. DAPI is used as a nuclear stain. Scale bar: 20 µm. e) Quantification of CD3^+^;CD8^+^ cells of IF stainings shown in d) (n=5). A whole slide scan of stained prostate tissue per mouse was analyzed. The percentage of positive cells in the prostate epithelium relative to *Pten*^peΔ/Δ^ was calculated. f) Representative pictures of immunohistochemistry (IHC) staining of NimpR14 (higher panel) and F4/80 (lower panel) of mouse prostates with indicated genotypes. Scale bar: 40 µm. g-h) Quantification of NimpR14^+^ (g) and F4/80^+^ (h) cells in the prostate epithelium of IHC stainings shown in f) (n≥6). A whole slide scan of stained prostate tissue per mouse was analyzed. The percentage of positive cells in the prostate epithelium relative to *Pten*^peΔ/Δ^ was calculated. i) Representative pictures of H&E and immunohistochemistry (IHC) staining of CD3, NimpR14 and F4/80 (in presented order) of *Pten*^peΔ/Δ^, and *Pten*^peΔ/Δ^;Stat3^peΔ/Δ^ prostates. Scale bar: 40 µm. j) Semi-quantitative analysis of CD3, NimpR14 and F4/80 IHC stainings shown in i) (n≥3). (c,e,g-h,j) Individual biological replicates are shown (c,e,g-h). Data are plotted as the means±SD and p-values were determined by unpaired two-tailed Student’s t-tests (c,e,g-h) or Mann-Whitney test (j).

A more detailed characterization showed that B-cells (CD79a) were not involved in the immune cell infiltration (data not shown). Importantly, CD3^+^ cells were significantly increased in the prostate epithelium of *Pten*^peΔ/Δ^;*L-gp130*^peKI/KI^ mice (**Supplementary Fig. 5d-e**), whereas, in the adjacent stroma, the CD3^+^ cell levels did not exhibit a significant difference compared to *Pten*^peΔ/Δ^ mice (**Supplementary Fig. 5f**). Notably, a substantial proportion of these cells were CD3^+^;CD8^+^ positive (**Fig. 5d** and **Supplementary Fig. 5g-h**), which are considered major drivers of anti-tumor immunity^49^. We also noted a significant difference in the ability of CD3^+^;CD8^+^ cells to migrate into the epithelium between *Pten*^peΔ/Δ^ and *Pten*^peΔ/Δ^;*L-gp130*^peKI/KI^ mice (**Fig. 5e**), whereas their proportion in the stroma was the same in both genotypes (**Supplementary Fig. 5i**). Concurrently, our findings indicated that CD3^+^;CD4^+^ T-cells did not play a significant role in anti-tumor infiltration in our mouse model (**Supplementary Fig. 5j-l**). Next, we examined neutrophils and macrophages to understand their potential contributions to the immune response within the prostate epithelium and the TME. IHC stainings for the neutrophil marker NimpR14 and macrophage marker F4/80 revealed a significant increase in the epithelial fraction of *Pten*^peΔ/Δ^;*L-gp130*^peKI/KI^ compared to *Pten*^peΔ/Δ^ mice (**Fig. 5f-h**), but no change in stroma or in the comparison of wild type and *L-gp130*^peKI/KI^ mice (**Supplementary Fig. 5m-r**), reflecting innate immune cell tumor infiltration upon constitutive GP130 signaling activation in *Pten*^peΔ/Δ^;*L-gp130*^peKI/KI^ mice. In agreement with these data, selected adaptive and innate immune system-related gene sets associated with chemotaxis, migration, regulation, and activation of immune cells were significantly upregulated when comparing *Pten*^peΔ/Δ^;*L-gp130*^peKI/KI^ with *Pten*^peΔ/Δ^ samples (**Supplementary Fig. 5s**), further substantiating the importance of infiltrating immune cells, specifically T-cells, neutrophils and macrophages, in our mouse model of PCa.

Given the importance of inflammatory cytokines in regulating the recruitment and activation of T-cells, neutrophils, and macrophages, we screened the significantly deregulated HALLMARK gene sets for related genes sets. Indeed, the gene set “Inflammatory response” and signaling of the effector molecules IFNy and TNFα, which are secreted by cytotoxic T-cells and affect tumor cells^50,51^, were significantly upregulated in our RNA-Seq data set (**Fig. 4a**). In order to delineate the cytokine profile more comprehensively, we analyzed serum samples obtained from the PCa mouse model. We specifically assessed the expression levels of various cytokines, chemokines, and receptors, including VEGF, CCL5, TNFα, IL-1α, IL-2R, IL-12p70, CXCL1, CXCL5, CXCL10, CD27, G-CSF. The multiplex immunobead assay analysis revealed a significant alteration in the cytokine profile, characterized by a significant upregulation of the expression of inflammatory cytokines in the serum of *Pten*^peΔ/Δ^;*L-gp130*^peKI/KI^ mice compared to the *Pten*^peΔ/Δ^ group (**Supplementary Fig. 5t**). Taken together, these findings provide evidence that constitutively active GP130 signaling in prostate epithelial cells promotes the recruitment of T-cells, neutrophils, and macrophages and reshapes the TME towards higher infiltration susceptibility.

To provide mechanistic evidence that these alterations depend on STAT3 signaling, we utilized a previously established PCa mouse model featuring *Pten*^peΔ/Δ^ with an additional prostate epithelium-specific deletion of *Stat3* (*Pten*^peΔ/Δ^;*Stat3*^peΔ/Δ^). These mice exhibit rapid tumor proliferation, metastasis, and an early death, in contrast to the slow, localized tumor progression seen in *Pten*^peΔ/Δ^ mice^9^. Interestingly, several immune response-related pathways are downregulated in *Pten*^peΔ/Δ^;*Stat3*^peΔ/Δ^ compared to *Pten*^peΔ/Δ^ prostates^40^. Consistent with these findings and our data in *Pten*^peΔ/Δ^;*L-gp130*^peKI/KI^ mice, *Pten*^peΔ/Δ^;*Stat3*^peΔ/Δ^ prostates showed no increase in the infiltration of CD3^+^ T-cells and F4/80^+^ macrophages, and a significant decrease in NimpR14^+^ neutrophils compared to *Pten*^peΔ/Δ^ mice (**Fig. 5i-j**) highlighting the importance of STAT3 in immune cell infiltration in *Pten*-deficient PCa mice with concomitant active GP130 signaling.

### GP130 signaling in PCa patients promotes STAT3 activation, senescence upregulation, elevated immune scores, and T-cell mediated cytotoxicity

To address the human relevance of our findings concerning the involvement of senescence and anti-tumor immunity in the proposed tumor-suppressive role of GP130/STAT3 signaling, we refined our analysis of the TCGA-PRAD patient data set by distinguishing *GP130*^high^ and *GP130*^low^ groups based on *GP130* mRNA expression levels (**Fig. 3b**). The fGSEA analysis of HALLMARK gene sets revealed that „IL-6/JAK/STAT3 signaling” was upregulated in *GP130*^high^ compared with *GP130^low^* patients (**Fig. 6a**), as evidenced by increased STAT3 target genes expression (**Supplementary Fig. 6a**). The observed downregulation of „Oxidative phosphorylation” is in accordance with the inverse association with STAT3^40^ and the corresponding KEGG gene set (**Supplementary Fig. 6b**).

**Fig. 6:**
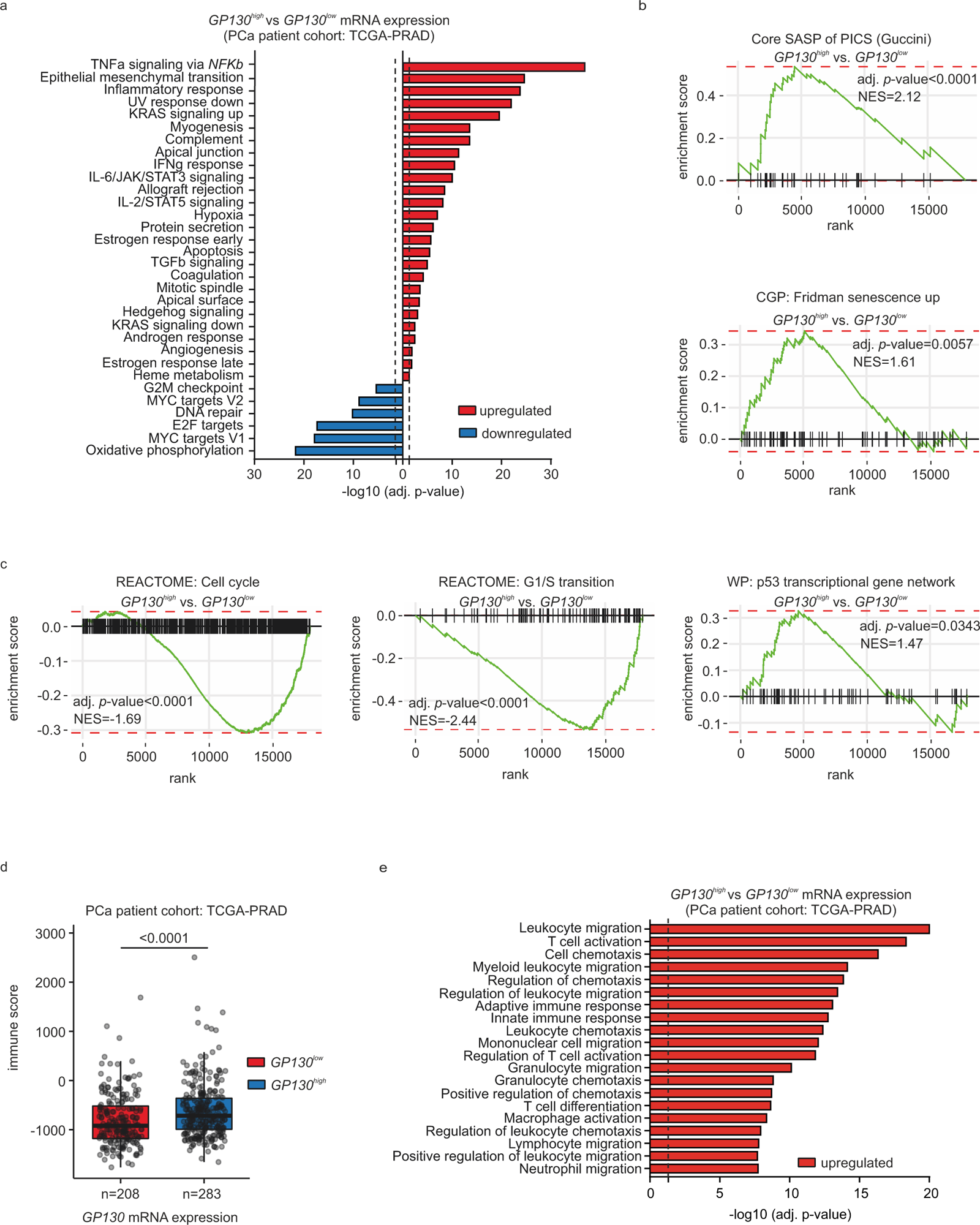
GP130 signaling in PCa patients leads to active STAT3 signaling, upregulation of senescence and higher immune score and T-cell mediated cytotoxicity. a) Fast pre-ranked gene set enrichment analysis (fGSEA) of significantly enriched HALLMARK gene sets with genes regulated in *GP130*^high^ compared to *GP130*^low^ expressing patients from the TCGA-PRAD data set. Dotted line: adj. p-value (-log10(0.05)), blue: downregulated, red: upregulated; b) Fast pre-ranked gene set enrichment analysis (fGSEA) of the previously described core SASP gene signature upon PICS “Core SASP of PICS (Guccini)” (upper panel) and the curated gene set, class chemical and genetic perturbations (CGP) “CGP: Fridman senescence up” (lower panel) with genes regulated in *GP130*^high^ compared to *GP130*^low^ expressing patients from the TCGA-PRAD data set. Genes sorted based on their Wald statistics are represented as vertical lines on the x-axis. NES: normalized enrichment score. c) Fast pre-ranked gene set enrichment analysis (fGSEA) of WikiPathways (WP) gene sets “REACTOME: Cell cycle”, “REACTOME: G1/S transition” and “WP: p53 transcriptional gene network” with genes regulated in *GP130*^high^ compared to *GP130*^low^ expressing patients from the TCGA-PRAD data set. Genes sorted based on their Wald statistics are represented as vertical lines on the x-axis. NES: normalized enrichment score. d) Immune score from the ESTIMATE method for *GP130*^low^ (red, n=208) and *GP130*^high^ (blue, n=283) patients from the TCGA-PRAD data set, compared with Mann-Whitney test. e) Fast pre-ranked gene set enrichment analysis (fGSEA) of the top 20 T-cell-, neutrophil-, and macrophage-associated Biological Processes from Gene Ontology pathways (GO-BP) gene sets with genes significantly regulated in *GP130*^high^ compared to *GP130*^low^ expressing patients from the TCGA-PRAD data set. Dotted line: adj. p-value (-log10(0.05)), red: upregulated;

As we depicted alterations in senescence and cell cycle regulators in our *in vivo* mouse model, we performed fGSEA excluding any patients with *TP53* mutations (**Supplementary Table 1**). Analysis of senescence-related gene sets (previously published “Core SASP of PICS (Guccini)“^47^ and “Fridman senescence up” taken from curated gene sets, class chemical and genetic perturbations (CGP)) revealed their significant upregulation in *GP130*^high^ PCa patients (**Fig. 6b**), providing a possible explanation for their improved survival (**Fig. 3b**). These patients also exhibited downregulation of cell cycle gene sets (“REACTOME: Cell cycle” and “REACTOME: G1/S transition”) and upregulation of p53 signaling (“WikiPathways (WP): p53 transcriptional gene network”) (**Fig. 6c**).

Using ESTIMATE (Estimation of STromal and Immune cells in MAlignant Tumor tissues using Expression data), a tool for predicting tumor purity, and the presence of infiltrating stromal/immune cells in tumor tissues based on gene expression data^52^, we confirmed that the majority of PCa patients can be considered immune-cold due to their low immune scores (**Supplementary Fig. 6c**). Notably, higher immune scores have been associated with longer survival rates in PCa patients^53^. In our patient cohort, *GP130*^high^ patients showed significantly higher immune scores compared to *GP130*^low^ patients (**Fig. 6d**), correlating with the upregulation of immune response-related gene sets (**Fig. 6a**). Furthermore, the top 20 GO-BP gene sets associated with T-cell activation and cytotoxicity, neutrophils and macrophages were upregulated in *GP130*^high^ compared to *GP130*^low^ expressing patients from the TCGA-PRAD data set, emphasizing the relevance of T-cell, neutrophil and macrophage mediated tumor-defense in PCa patients with high *GP130* expression (**Fig. 6e**). In summary, our data reveal that PCa patients with high *GP130* expression exhibit increased senescence, reduced cell cycle activity, and enhanced immune cell infiltration, which likely contribute to their improved survival outcomes.

## Discussion

In this study, we show that in a *Pten*-deficient PCa mouse model engineered to constitutively activate GP130 signaling, STAT3 activation was increased, STAT3 target gene signature was amplified and PCa tumor growth was significantly reduced compared to mice only deficient in *Pten*. The proposed roles of active GP130 signaling in PCa observed in this study are summarized in **Figure 7**. We found that enhanced STAT3 signaling was associated with more pronounced p19^ARF^/p53 mediated cellular senescence in the tumor tissue. These findings complement previous data showing that inducing a *Stat3* knock out (KO) in PCa mice resulted in larger tumor sizes mediated by loss of senescence^8,9^. The STAT3 signaling axis thus appears to regulate tumor growth in PCa by primarily inhibiting tumor progression rather than initiation. This is evidenced by the majority of *Pten*^peΔ/Δ^;*L-gp130*^peKI/KI^ mice displaying PINs and not PCa, as was predominantly found in *Pten*^peΔ/Δ^ mice.

**Fig. 7:**
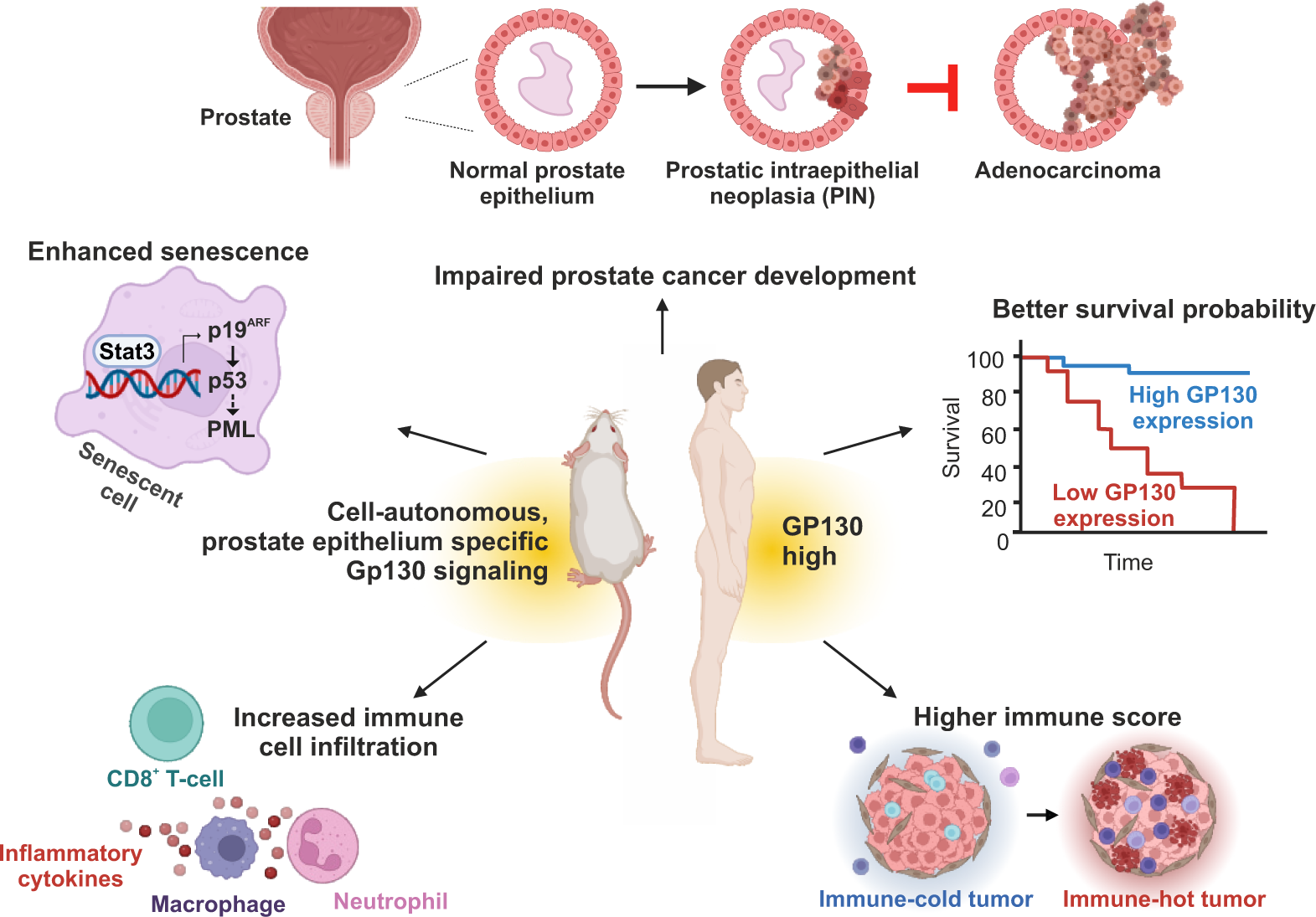
Proposed roles of active GP130 signaling in prostate tumorigenesis. Using the genetic mouse model, we showed that cell-autonomous, prostate epithelium-specific and constitutively active GP130 signaling reduces *Pten*-deficient tumor growth, enhances the STAT3/p19^ARF^/p53-driven senescence and recruits tumor-infiltrating immune cells (T-cells, neutrophils, and macrophages). In human PCa, high *GP130* expression causes active STAT3 signaling, correlates with better survival and is associated with higher level of immune infiltrates.

The data presented here provide substantial validation and significantly broaden the scope of our previously posited hypotheses^9^, that 1) the STAT3/p19^ARF^ axis acts as a safeguard mechanism against malignant progression in PCa, 2) expression levels of constituents of the GP130/STAT3 signaling axis could act as key markers to stratify PCa cases into low- and high-risk groups, and 3) strategies aimed at manipulating this signaling pathway could serve as a novel therapeutic approach for PCa treatment.

The tumor-suppressive role of STAT3 signaling in PCa contrasts with the oncogenic function in numerous cancers, where it is often hyperactivated^5,54,55^. However, accumulating evidence suggests that it may also function as tumor suppressor depending on signaling context and tumor type^56,57^. For example, *Stat3* deletion resulted in increased astrocyte tumor formation in SCID mice in the absence but not in the presence of PTEN^58^. Additionally, in a mouse model of colorectal cancer crossed with *Stat3* conditional KO mice, *Stat3* KO in intestinal cells revealed an oncogenic role, whereas a KO during tumor progression enhanced tumor invasiveness, reflecting a tumor suppressive role^59^. In a more specific example, a mouse model of drug-induced cancer demonstrated that STAT3 expression appeared to suppress tumor formation in the presence of a toxicant causing chronic liver injury, inflammation, and fibrosis, whereas it enhanced tumor formation induced by a DNA damaging agent^60^. Further examples have been reported for lung cancer, thyroid cancer and head and neck squamous cell cancers^56^, additionally supporting the notion that STAT3 signaling has a dual role, rather than a strictly oncogenic one.

As a possible molecular mechanism underlying this ambiguous behavior, activity of the STAT3β isoform has been suggested. This isoform lacks the C-terminal transactivation domain and was shown to inhibit proliferation and stimulate cell death, possibly through heterodimerizing with STAT3α, thereby preventing it from activating its target genes^57^. In contrast, our data suggest that a different mechanism mediates the tumor suppressive activity of STAT3 signaling in prostate cells with constitutively activated GP130. This assumption is based on the observed clear upregulation of transcription in both the absence and presence of PTEN (*Pten*^peΔ/Δ^;*L-gp130*^peKI/KI^ and *L-gp130*^peKI/KI^ mice), as determined from differential gene expression analyses. Another aspect where context dependency seems to be decisive is senescence and its associated SASP. Both have nuanced roles in PCa, with outcomes potentially shaped by context and genetic backgrounds^61^. Examining the transcriptome upon constitutively active GP130 signaling in greater detail, we identified a significant upregulation of the senescence-associated p19^ARF^/p53 pathway and PICS. Targeting senescence in this context has been suggested to potentially hold significant promise in cancer therapy^45,62,63^. However, a tumor-promoting effect linked to increased SASP in a PCa mouse model with additional KO of *Stat3* has been reported^64^. One possible factor contributing to this divergence in outcomes might be influenced by the specific *Stat3* KO approach that targets only the tyrosine phosphorylation site of *Stat3* and not the DNA-binding domain^65^. Variations in STAT3 expression, such as dominant-negative STAT3, can have profound implications on disease outcomes^66^. Our studies, spanning four independent mouse model systems addressing the IL-6/GP130/JAK2/STAT3 signaling axis, consistently indicate a tumor-suppressive effect^8,9^. Specifically, our genetic PCa mouse model with KO of *Stat3*, in which the DNA-binding domain of *Stat3* is targeted^67^, leads to aggressive PCa growth. Additionally, we have previously shown that the KO of *Il6*, the activator of the GP130/STAT3 signaling, enhances PCa development^9^. By using an independent mouse model, our current study additionally emphasizes a tumor-suppressive role for constitutively active GP130/STAT3 signaling and its associated elevated SASP. This divergence in findings highlights the need to consider context- and patient-specific factors, further challenging the current discussion on the therapeutic advantages or hazards of IL-6/STAT3 inhibition.

To assess the potential clinical relevance of our murine findings, we analyzed several PCa patient cohorts. We discovered that in the overall patient population *GP130* expression was significantly reduced in the prostate tumor tissue compared to surrounding non-cancerous tissue. However, when we separated these patients into a high and a low *GP130* expressing group, we detected that higher *GP130* expression correlated with higher *STAT3* expression and, more importantly, with prolonged recurrence-free survival of these patients. Therefore, based on the hypothesis that enhanced GP130 signaling, as observed in our mouse model, helps restrain cancer progression, we propose that *GP130* expression levels, possibly along with *STAT3* and *ARF* levels, could serve as valuable marker for identifying low- and high-risk PCa groups. This distinction could help prevent overtreatment and unnecessary reductions in quality of life for PCa patients^68^.

A further important potential lead for future treatment of PCa patients derived from our study is the observation that enhanced GP130 signaling was associated with high-grade immune cell infiltration at the tumor site. This infiltration included CD3^+^;CD8^+^ T-cells, neutrophils, and macrophages that are considered major drivers of anti-tumor immunity^49,69^, and was accompanied by an upregulation of adaptive and innate immune system-related gene sets. In general, PCa cells and those comprising its microenvironment are known to express and secrete molecules mediating immunosuppression, rendering PCa immune-cold^22^ and thus not a good target for otherwise highly efficient immune-based therapies^70^. PCa is also associated with low mutational burden and low immunogenicity^71,72^, and thus little responsiveness to therapies based on immune checkpoint inhibitors^73^. Consequently, the potential to enhance immunogenicity by manipulating GP130 signaling could present a novel therapeutic approach in PCa therapy.

Our analysis of human patient data sets supports this idea, confirming that, while most PCa patients examined must be regarded as immune-cold (as determined by the ESTIMATE tool^52^) those with high *GP130* expression exhibited a better immune score. These patients also showed upregulated gene sets associated with T-cells, neutrophils, and macrophages and, noteworthy, increased senescence-related gene sets. As to the latter, it remains to be shown whether immune cell infiltration is the direct consequence of enhanced GP130 signaling and the associated senescence induction or its cause^61,74^. In either case, there is evidence suggesting that a higher number of tumor infiltrating lymphocytes in PCa is associated with better patient outcomes^75^. Considering the immense potential of novel approaches, such as the induction of synthetic cytokine signaling circuits allowing immune cells to overcome immunosuppressive microenvironments and infiltrate immune-excluded solid tumors^76^, it appears conceivable that strategies mediating prostate-specific, active GP130 signaling have potential to effectively attack tumor cells.

Altogether, the present study reveals that increased GP130 signaling is linked with suppressed tumor growth and amplified STAT3 target gene signatures. Contrary to its oncogenic role in many cancers, GP130/STAT3 signaling demonstrated tumor-suppressive activity in the context of PCa, potentially through the upregulation of the senescence-associated p19^ARF^/p53 pathway. Additionally, elevated GP130 signaling in tumors was linked to increased immune cell infiltration, implying that enhancing GP130 signaling might be a promising therapeutic strategy for boosting anti-tumor immunity in PCa. Clinical analysis of PCa patients showed a positive correlation between high *GP130* expression and longer recurrence-free survival, suggesting *GP130* expression levels could serve as markers for risk stratification.

## Material and Methods

### Generation of transgenic mice

*Pten*^fl/fl27^, *L-gp130*^fl/fl4^ and PB-Cre4^26^ transgenic mice were maintained on a C57BL/6 and Sv/129 mixed genetic background. *Pten*^fl/fl^ mice and/or *L-gp130*^fl/fl^ mice were crossed with male PB-Cre4 transgenic mice to generate prostate-specific deletion of *Pten* and/or insertion of the *L-gp130* construct. DNA isolation was performed as previously described^29^. Mice were genotyped as previously described^4,25–27^. Mice were housed on a 12–12 light cycle and provided food and water ad libitum. For all experiments, 19 weeks old male mice were used. Genotyping primer sequences and protocols are listed in **Supplementary Table 2-3**. Formalin-fixed paraffin-embedded (FFPE) prostate tissue from *Pten*^peΔ/Δ^;*Stat3*^peΔ/Δ^ and respective *Pten*^peΔ/Δ^ control mice were provided by ^9^.

### Immunohistochemistry and hematoxylin & eosin stains

Immunohistochemistry (IHC) and hematoxylin & eosin stains (H&E) were performed with FFPE prostate tissue using standard protocols and antibodies listed in **Supplementary Table 4**. Representative pictures for figures were exported from whole slide scans using the snapshot function of CaseViewer (Build 2.4.0.119028).

### Immunofluorescence staining

Frozen tissue sections were fixed with 4% Formol for 15 min at room temperature. After washing with PBS, cells were blocked with 2% BSA in PBS prior to overnight primary antibody incubation at 4°C (**Supplementary Table 4**). Secondary antibody incubation (Alexa Fluor 594 anti-rabbit, Invitrogen #A11037, 1:500) was done in 2% BSA in PBS for 1 h at room temperature. Cells were counterstained with DAPI (nuclear stain) in PBS and mounted with Aqua-Poly/Mount medium (18606-5, Polysciences). Representative pictures for figures were exported from whole slide scans using the snapshot function of CaseViewer (Build 2.4.0.119028).

### Multiplex immunohistochemistry

Mouse prostate samples were stained with multiplex immunohistochemistry and analyzed by multispectral imaging. A panel of the following fluorescent markers plus DAPI as a nuclear stain were used to detect the following epitopes: CD3, CD4, CD8, CD45, pan Cytokeratin (**Supplementary Table 4**). Staining was performed with the autostainer system Bond RX (Leica Biosystems Inc., Vienna, Austria). The slides were scanned with the Vectra® 3 (Akoya Biosystems, Marlborough, USA; software version 3.0.7) microscope. Whole-slide scans were taken at 4x magnification to define regions of interest (whole tissue area) to be scanned in higher resolution using Phenochart software, version 1.0.12. Multispectral images of the defined areas (whole tissue) were then recorded at 20x magnification, resulting into one image color channel for each stained antibody. Images were processed with inForm software (Akoya Biosystems, Marlborough, USA; software version 2.4.10), including spectral unmixing and removal of autofluorescence. Resulting multispectral images were evaluated using HALO® Image Analysis Platform (Indica Labs, Albuquerque, NM, US). Single recorded images at 20x magnification were stitched together into a continuous field of view of the whole tissue. Individual cells were identified using the DAPI nucleus staining by setting a threshold for nucleus size, roundness, and signal intensity. For the fluorescent labelled markers, positivity thresholds were set according to the staining intensity.

### Histopathological analysis

For histopathological analysis, a whole slide scan of stained prostate tissue per mouse was analyzed. Stained slides were scanned with a PANNORAMIC Scan II from 3DHISTECH, using the following parameters: Objective type: 20x; Output resolution: 49x native; Multilayer mode: extended focus, 7 levels, step size 1 µm; Compression: JPG; Bit depth: 8-bit; stitching enabled; and saved as MRXS files. Quantitative analysis of immunohistochemistry stainings was performed with QuPath^77^ (version 0.3.2). First, regions of interest were annotated, excluding non-prostate tissue such as urethra, seminal vesicles, and ductus deferens. Cell detection was performed with the StarDist^78^ extension for the NimpR14 staining and the built-in watershed cell detection plugin for the other stainings. Parameters were chosen individually for each staining. Thereafter smoothed features were calculated with a FWHM radius of 25 µm. The tissue was then classified into epithelium and stroma using an object classifier, trained individually for each staining. For the evaluation of F4/80 and NimpR14, only the caudal prostate lobe was used due to its clearer morphological architecture. A threshold was set for the mean DAB optical density value, categorizing cells into positive or negative. If automated quantification was not possible for immunohistochemical stainings, semi-quantitative analysis was performed by a trained pathologist, who classified the level of expression as none, mild, moderate or marked for each tissue section. A similar approach was taken for the grading of immune cell infiltration, which was classified as low or high in H&E-stained sections. Analyses were performed blinded to genotype by a single investigator and evaluated by two independent pathologists with specific expertise in mouse models of PCa. Representative pictures for figures were exported from whole slide scan using the snapshot function of CaseViewer (Build 2.4.0.119028).

### Quantitative RT-PCR (qRT-PCR)

Mouse prostate tissue was treated with RNAlater Stabilization Solution (AM7020, Thermo Fisher Scientific, USA) according to the manufacturer’s instructions. RNA isolation was performed using TriReagent (T9424-100ML, Sigma-Aldrich) and ReliaPrep RNA Tissue Miniprep System (Z6111, Promega) according to the manufacturer’s instructions. DNase digestion was performed on a column. Conversion of one µg of RNA into cDNA was performed using the iScript cDNA Synthesis Kit (1708890; Bio-Rad Laboratories) and Master Cycler Pro Device (EPPE6324000.516, Eppendorf, DE). For RT-qPCR, 2xBiozym Blue ŚGreen qPCR Kit (331416, Biozym Scientific GmbH) was used and analysis was done on a ViiA 7 Real-Time PCR System (4453536, Thermo Fisher Scientific). mRNA levels were normalized to the geometric mean of Cyclophilin A and hypoxanthine guanine phosphoribosyl transferase and/or 18S. The sequences of primers used for amplification are listed in **Supplementary Table 2**.

### Protein isolation and immunoblotting

SDS-PAGE and Western blotting were performed as previously described^79^. Whole prostate protein lysates were extracted from snap frozen prostate samples and 20-40 μg of protein lysate were used. Chemiluminescent visualization was performed with a ChemiDoc™ Imaging System (Bio-Rad) after incubation of the membranes with Clarity Western ECL reagent (Bio-Rad). Quantifications were performed with Image Lab software (Bio-Rad). Samples were normalized to the indicated loading controls. Applied antibodies are listed in **Supplementary Table 4**.

### Multiplex immunobead cytokine assay

Mouse serum samples were collected and analyzed using the ProcartaPlex antibody-based, magnetic bead reagent assay panels. This approach utilizes Luminex xMAP technology and the associated instrument platform for multiplex protein quantitation. We simultaneously determined the concentrations of cytokines in supernatant samples using a customized 28-plex immunoassay kit (ProcartaPlex Mouse 28-plex, ThermoFisher Scientific), which employs magnetic beads for detection. The samples, which were undiluted and stored frozen -80°C, were processed upon thawing in a 96-well plate following the manufacturer’s instructions. We generated standard curves for each analyte by measuring individual standards in duplicate. These measurements were referenced against concentrations supplied by the manufacturer. The assays were conducted using a calibrated Bio-Plex 200 system (Bio-Rad), and data analysis was performed with the Bio-Plex Manager software, version 6.1 (Bio-Rad). We calculated the cytokine concentrations from the standard curves employing five-parameter logistic (5PL) regression curve fitting. The fluorescent intensity measurement of 11 out of 28 cytokines/chemokines were within the standard curves highest and lowest value point (CXCL5, IL-1α, VEGF, G-CSF, IL-12p70, CXCL1, IL-2R, CD27, CXCL10, CCL5 and TNFα).

### RNA sequencing (RNA-Seq)

RNA-Seq sample and library preparation was performed as described in ^29^. Briefly, single cell suspension of mouse prostate tissue of wild type, *L-gp130*^peKI/KI^, *Pten*^peΔ/Δ^ and *Pten*^peΔ/Δ^;*L-gp130*^peKI/KI^ mice was done as previously described^80^ and magnetic cell separation (MagniSort technology, Thermo Fisher) was performed for EpCAM positive fraction using anti-mouse CD326 (EpCAM) Biotin antibody (13-5791-82, eBioscience). For higher RNA output, three wild type and three *L-gp130*^peKI/KI^ mouse prostates, respectively, were pooled to generate one sample. Per genotype n≥5 samples were analyzed by RNA-Seq. RNA isolation was performed using TriReagent (T9424-100ML, Sigma-Aldrich) and ReliaPrep RNA Tissue Miniprep System (Z6111, Promega) according to the manufacturer’s instructions. Library preparation was performed using NEBNext Ultra II Directional RNA Library Prep Kit for Illumina (E7760, New England Biolabs) according to manufacturer’s instructions in combination with a poly(A) mRNA magnetic isolation module (E7490) and multiplex oligos for Illumina (E7600). Libraries were amplified with 11 PCR cycles and the library size was analyzed by Agilent Tape Station (G2938-90014, Agilent Technologies).

### RNA-Seq data analysis

RNA sequencing and bioinformatic analysis of mouse prostate samples up to the differential expression was performed by Core Facility Bioinformatics of CEITEC Masaryk University as previously described^29^.

### Fast pre-ranked gene set enrichment analysis (fGSEA)

fGSEA analysis was performed in R (version 4.2.0). Gene sets used for the fGSEA analysis were derived from MSigDB^30^ (version 7.5.1; collections: HALLMARK pathway database^31^, REACTOME^81^, Wikipathways^82^, Gene Ontologies (GO)^83,84^, gene set: Fridman_senescence_up (M9143)) through the msigdbr^85^ (version 7.5.1) R package or from previously published works („core SAPS of PICS“^47^, “STAT3 targets (Swoboda)”^32^, “STAT3 targets (Azare)”^33^, “STAT3 targets (Carpenter)”^34^). All expressed genes within an experiment were extracted and sorted based on their Wald statistics obtained during the differential expression analysis performed with DESeq2. Sorted list of genes was used as an input for fGSEA^86^ (version 1.22.0) R package. Human or mouse gene symbols present in custom gene sets were converted to orthologous mouse or human genes, respectively, using R package biomaRt^87^ (version 2.52.0) when necessary.

### TCGA data analysis

Clinical data for the TCGA-PRAD cohort (https://portal.gdc.cancer.gov/projects/TCGA-PRAD)^35^, including disease-free survival, were downloaded from the cBioportal^88,89^ database. Raw expression counts were downloaded from TCGA with the TCGAbiolinks R package (version 2.25.3). Patients with mutation in TP53 gene (**Supplementary Table 1**) were removed from subsequent analysis for Fig. 6b-c. Raw counts were transformed with the Variance-stabilizing transformation. Survival analysis of TCGA-PRAD cohort was performed with the survminer R package^90^ (version 0.4.9). Patients were divided into *GP130*^high/low^ expression groups based on the maximally selected rank statistics which provides a single value cutpoint that corresponds to the most significant relation with disease-free survival. Differential expression analysis between *GP130*^high^ and *GP130*^low^ group was performed with DESeq2 (version 1.36.0). Alteration data including frequency, co-occurrence of specific mutations and correlation analysis were obtained from the cBioportal^88,89^ database. Immune scores for TCGA-PRAD cohort from the ESTIMATE method^52^ were downloaded from https://bioinformatics.mdanderson.org/estimate/. Difference between *GP130*^high^ and *GP130*^low^ based on the immune score was estimated with Mann-Whitney test.

### MSKCC data analysis

Survival analysis and risk assessment of the publicly available data set Taylor MSKCC Prostate GSE21032^38^ were done using the SurvExpress online tool^37^. The prognostic index of *GP130* was estimated by fitting a Cox proportional hazards model. Risk groups were separated by ranking samples by their prognostic index median. They were analyzed by a concurrent Cox model and used for Kaplan-Meier plots and log-rank tests. Biochemical recurrence was determined by an increase of >0.2 ng/ml PSA in serum. Correlation analysis was done using the cBioportal^88,89^ database.

### Oncomine database analysis

Gene expression data for *GP130* (*IL6ST*) were extracted from the following data sets using the Oncomine™ Research Premium Edition database (Thermo Fisher, Ann Arbor, MI)^36^: Arredouani Prostate (reporter: 204863_s_at), Lapointe Prostate (reporter: IMAGE:2018581), Wallace Prostate (reporter: 204863_s_at), Varambally Prostate (reporter: 204863_s_at and 204864_s_at), Grasso Prostate (reporter: A_32_P140656), Holzbeierlein Prostate (reporter: 35842_at), Taylor Prostate 3 (reporter: 6733), Glinsky Prostate (reporter: 212196_at) and Best Prostate 2 (reporter: 212196_at). Statistics are reported as they appear in the Oncomine database.

### Statistical analysis

Significant differences between two groups were determined using a two-tailed, unpaired t-test (parametric) or Mann-Whitney test (non-parametric). Significant differences between more than two groups were determined using One-way ANOVA with Tukey’s multiple comparisons test (parametric). Significant outliers were identified by Grubbs’ test. p values of <0.05 were assigned significance. All values are given as means ± standard deviation (SD) and were analyzed and plotted by GraphPad Prism® (version 9.5.0, GraphPad Software, San Diego, CA). Numbers of biological replicates are stated in the respective figure legends.

### Ethics

Animal experiments and care were conducted in accordance with the guidelines of institutional authorities and approved by the Federal Ministry of Science, Research and Economy (BMWFW-66.009/0307-WF/V/3b/2017 and the associated amendments).

## Funding

This study was financially supported by grant Nr. 70112589 from the Deutsche Krebshilfe, Bonn, Germany to S.R.-J. L.K. acknowledges the support from MicroONE, a COMET Modul under the lead of CBmed GmbH, which is funded by the federal ministries BMK and BMDW, the provinces of Styria and Vienna, and managed by the Austrian Research Promotion Agency (FFG) within the COMET-Competence Centers for Excellent Technologies-program. Financial support was also received from the Austrian Federal Ministry of Science, Research and Economy, the National Foundation for Research, Technology and Development, the Christian Doppler Research Association and Siemens Healthineers. L.K. was also supported by a European Union Horizon 2020 Marie Sklodowska-Curie Doctoral Network grants (ALKATRAS, n. 675712; FANTOM, n. P101072735 and eRaDicate, n. 101119427) as well as BM Fonds (n. 15142), the Margaretha Hehberger Stiftung (n. 15142), the Christian Doppler Lab for Applied Metabolomics (CDL-AM), and the Austrian Science Fund (grants FWF: P26011, P29251, P 34781 as well as the International PhD Program in Translational Oncology IPPTO 59.doc.funds). P.W. was supported by the Austrian Science Fund (FWF - W1241). Additionally, this research was funded by the Vienna Science and Technology Fund (WWTF), grant number LS19-018. L.K. is a member of the European Research Initiative for ALK-Related Malignancies (www.erialcl.net). The National Institute for Cancer Research (Programme EXCELES, No. LX22NPO5102) funded by Next Generation EU is gratefully acknowledged for funding.

## Acknowledgements

We acknowledge the Core Facility Bioinformatics supported by the NCMG research infrastructure (LM2023067 funded by MEYS CR) for their support with the bioinformatic analysis of the RNA sequencing data presented in this paper. This research was supported using resources of the VetCore Facility (VetImaging) of the University of Veterinary Medicine Vienna. FFPE prostate tissue from *Pten*^peΔ/Δ^;*Stat3*^peΔ/Δ^ and respective *Pten*^peΔ/Δ^ control mice were obtained from ^9^. The authors are grateful to Dr. Natalie Bordag (Medical University of Graz) for providing help with R programming and bioinformatic analysis, and to Anton Jäger (Medical University of Vienna) for his assistance in generating macroscopic pictures and graphical illustrations. Scientific coaching to C.S. was provided by Gerhard Krumschnabel (medical-writing.at) and was funded by University of Veterinary Medicine Vienna. We acknowledge the use of BioRender.com for creating Figure 1a, Figure 4j and Figure 7.

## Author contributions

Conceptualization and design: C.S., S.R.-J., L.K.

Development of methodology: C.S., S.R.-J.,T.L., M.R., K.T., M.O., F.S., S.H., S.L.

Acquisition of data: C.S., T.L., M.R., K.T., D.L., M.S., R.Z., H.A.N., S.D., M.O., F.S.

Technical assistance: M.S., P.K., J.Y., B.T., M.T., N.S.H., S.S., S.T.

Project administration: C.S., S.R.-J., L.K.

Formal analysis: C.S., M.R., K.T., V.H., F.S.

Analysis and interpretation of data: C.S., T.L., H.A.N., T.R., F.S., S.H., S.L.

Resources: C.S., V.B., J.L.P., S.P., P.W., S.R.-J., L.K.

Study supervision: S.R.-J., L.K.

Funding acquisition: S.R.-J., L.K.

Writing - original draft: C.S.

Figure preparation: C.S., M.R., M.S., F.S.

Writing - review & editing: C.S., F.S., S.R.-J., L.K.

All authors read and approved the final manuscript.

## Competing interests

The other authors declare no competing interests.

## Abbreviations

adj. p-value: adjusted p-value
ARF: alternative reading frame
DE: differentially expressed
EpCAM: epithelial cell adhesion molecule
ERK: Extracellular signal-regulated kinase
ESTIMATE: Estimation of STromal and Immune cells in MAlignant Tumor tissues using Expression data
fGSEA: fast pre-ranked gene set enrichment analysis
fl: floxed
GP130: Glycoprotein 130 kDa
GOBP: Gene Ontology Biological Process
H&E: hematoxylin & eosin
IF: immunofluorescence
IHC: immunohistochemistry
IL-6: Interleukin-6
JAK: Janus kinase
KEGG: Kyoto Encyclopedia of Genes and Genomes
KI: knock in
KO: knock out
L-gp130: Leucine-gp130
MAPK: Mitogen-activated protein kinase
NES: normalized enrichment score
OIS: oncogene-induced senescence
p: phospho
PB: Probasin
PCa: prostate cancer
PCR: polymerase chain reaction
pe: prostate epithelium
PICS: PTEN-loss induced cellular senescence
PIN: prostatic intraepithelial neoplasia
PI3K: Phosphatidylinositol 3-kinase
PML: Promyelocytic leukemia protein
PTEN: Phosphatase and tensin homolog
RNA-Seq: RNA sequencing
SASP: senescence-associated secretory phenotype
SD: standard deviation
SHP2: Src homology 2 domain-containing tyrosine phosphatase-2
STAT3: Signal transducer and activator of transcription 3
t: total
TCGA-PRAD: The Cancer Genome Atlas-PRostate ADenocarcinoma
TMA: tissue microarray
TME: tumor microenvironment
WP: WikiPathways
ZSGreen: Zoanthus sp. green fluorescent protein

## Notes

### Competing Interest Statement

The authors have declared no competing interest.

