## Supplementary figures and tables for "Cell-autonomous GP130 activation suppresses prostate cancer development via STAT3/ARF/p53-driven senescence and confers an immune-active tumor microenvironment"

Supplementary Fig. 1

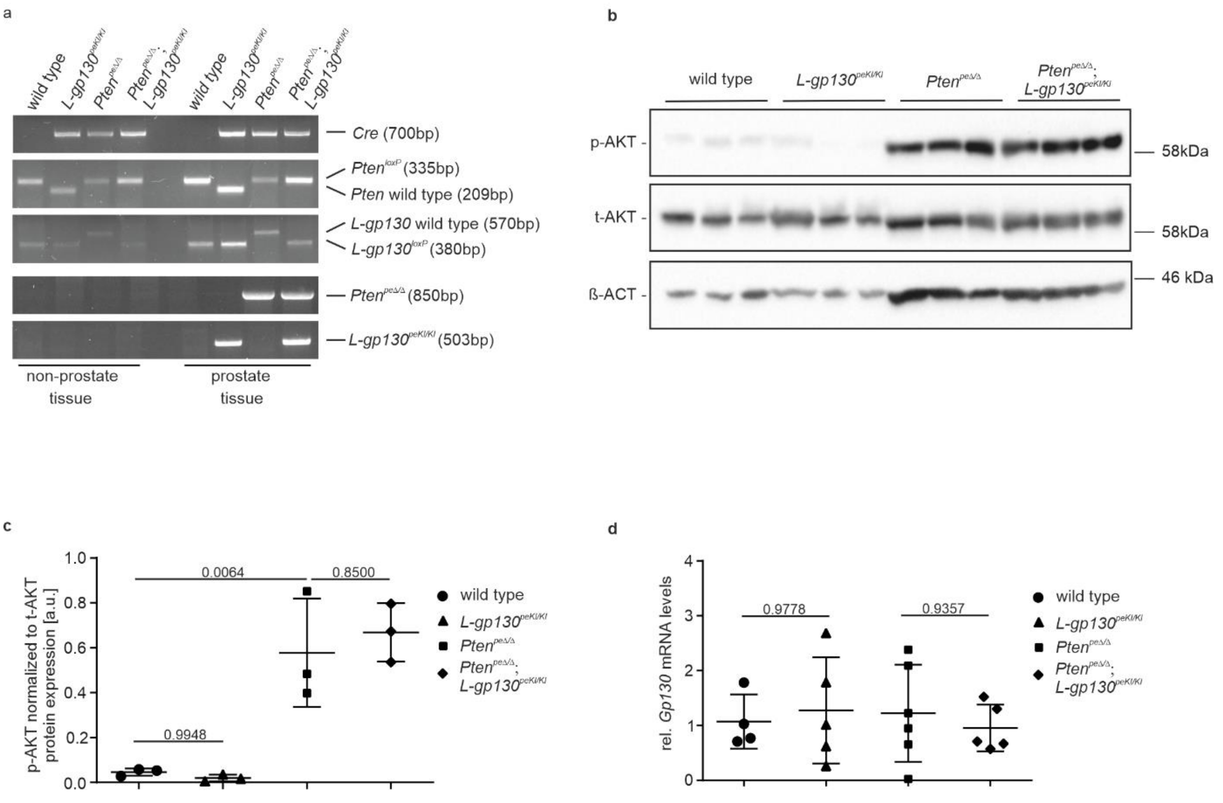


**Supplementary Fig. 1**

a) PCR-based analysis of the *Cre* transgene and non-recombined or recombined *Pten* and *L-gp130* alleles in genomic DNA purified from non-prostate tissue (left) and prostate tissue (right) from wild type (=littermates without *PB-Cre4*), *L-gp130*$^{peKI/KI}$, *Pten*$^{pe\Delta/\Delta}$, and *Pten*$^{pe\Delta/\Delta}$;*L-gp130*$^{peKI/KI}$ mice.

b) Representative Western blot analysis of prostate protein lysates for phospho-AKT (p-AKT) and total-AKT (t-AKT) expression in wild type, *L-gp130*$^{peKI/KI}$, *Pten*$^{pe\Delta/\Delta}$, and *Pten*$^{pe\Delta/\Delta}$;*L-gp130*$^{peKI/KI}$ mice (n=3). β-ACTIN (β-ACT) served as loading control.

c) Quantification of phospho-AKT (p-AKT) protein levels relative to total-AKT (t-AKT) protein levels from b).

d) Endogenous *Gp130* mRNA expression analysis by RT-PCR of wild type, *L-gp130*$^{peKI/KI}$, *Pten*$^{pe\Delta/\Delta}$, and *Pten*$^{pe\Delta/\Delta}$;*L-gp130*$^{peKI/KI}$ prostate tissue (n≥4).

(c-d) Individual biological replicates are shown. Data are plotted as the means±SD and p-values were determined by ordinary one-way ANOVA with Tukey's multiple comparisons.

**Supplementary Fig. 2**

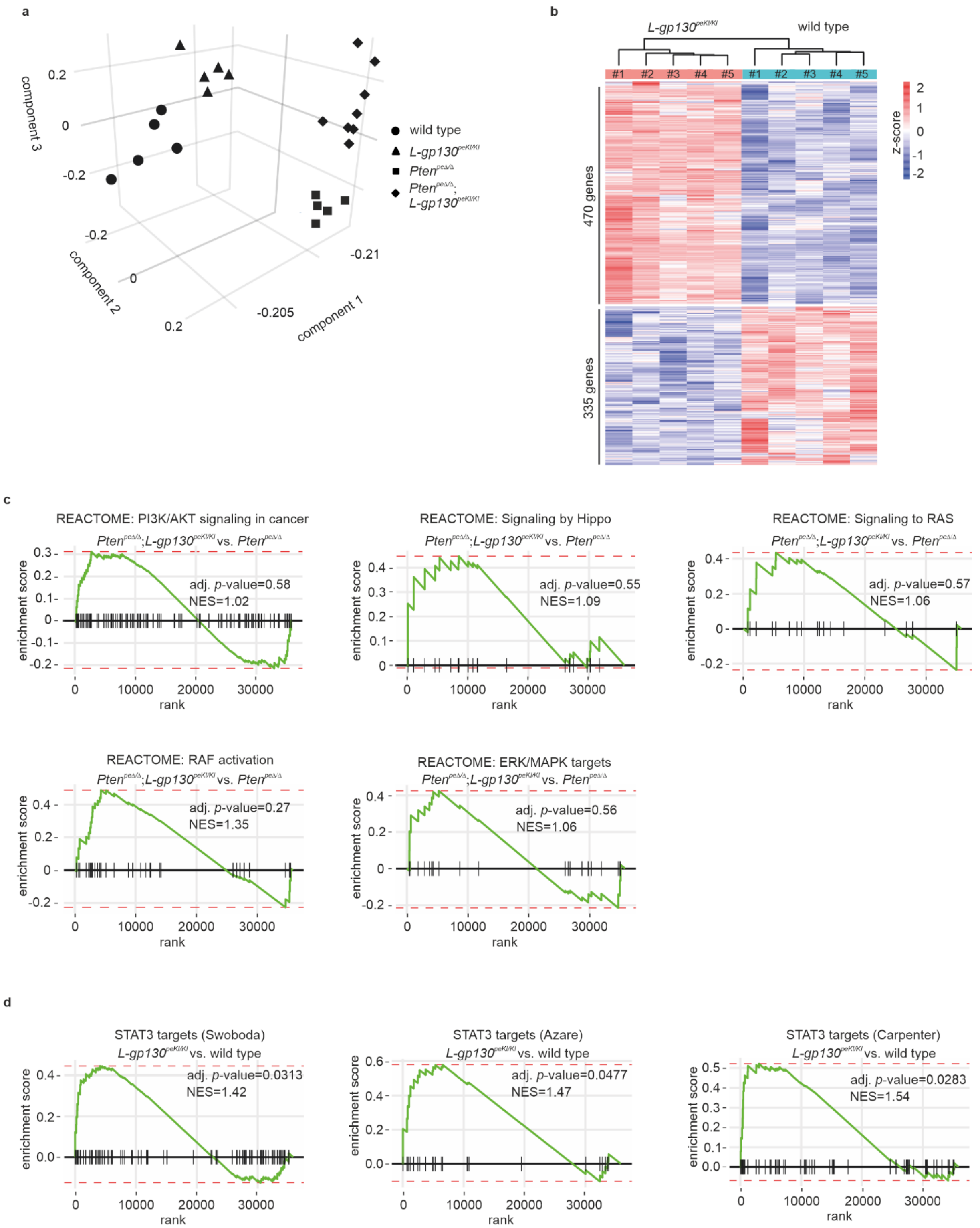

Supplementary Fig. 2 (continuation)

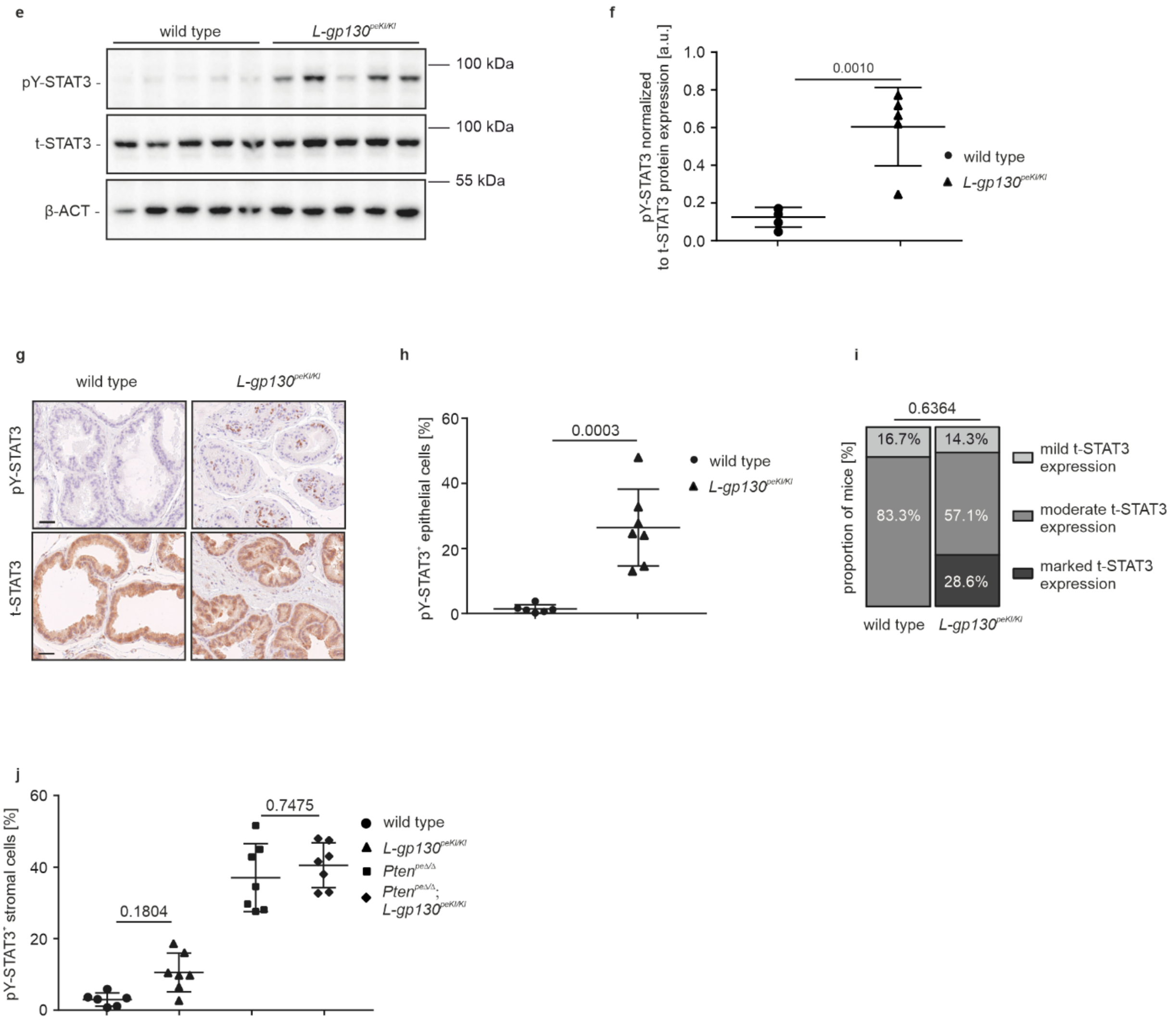


**Supplementary Fig. 2**

a) 3D-Principal component analysis based on mRNA gene expression of wild type, *L-gp130*$^{\mathrm{peKI/KI}}$, *Pten*$^{\mathrm{pe\Delta/\Delta}}$, and *Pten*$^{\mathrm{pe\Delta/\Delta}}$;*L-gp130*$^{\mathrm{peKI/KI}}$ prostate epithelial cells (n≥5).

b) Heatmap and number of differentially expressed genes (log2norm) based on adj. p-value ≤0.05 and fold change ≥2 cut-off values comparing *L-gp130*$^{\mathrm{peKI/KI}}$ and wild type prostate epithelial cells (n=5). blue: downregulated, red: upregulated.

c) Fast pre-ranked gene set enrichment analysis (fGSEA) of REACTOME gene sets (“PI3K/AKT signaling in cancer”, “Signaling by Hippo”, “Signaling to RAS”, “RAF activation”, “ERK/MAPK targets”) with genes regulated in *Pten*$^{\mathrm{pe\Delta/\Delta}}$;*L-gp130*$^{\mathrm{peKI/KI}}$ compared to *Pten*$^{\mathrm{pe\Delta/\Delta}}$ prostate epithelial cells. Genes sorted based on their Wald statistics are represented as vertical lines on the x-axis. NES: normalized enrichment score.

d) Fast pre-ranked gene set enrichment analysis (fGSEA) of three previously published STAT3 target signatures (“STAT3 targets (Swoboda)”, “STAT3 targets (Azare)”, “STAT3 targets (Carpenter)”) with genes regulated in *L-gp130*$^{\mathrm{peKI/KI}}$ compared to wild type prostate epithelial cells. Genes sorted based on their Wald statistics are represented as vertical lines on the x-axis. NES: normalized enrichment score.

e) Western Blot analysis of prostate protein lysates for phosphoTyrosine705-STAT3 (pY-STAT3) and total-STAT3 (t-STAT3) expression in wild type and *L-gp130*$^{peKI/KI}$ mice (n=5). β-ACTIN (β-ACT) served as loading control.

f) Quantification of pY-STAT3 protein levels relative to t-STAT3 protein levels shown in e).

g) Representative pictures of IHC staining of pY-STAT3 and t-STAT3 expression in prostate sections of wild type and *L-gp130*$^{peKI/KI}$ mice. Scale bar: 40 µm.

h-j) Quantitative analysis of STAT3 (positive epithelial cells in h, positive stromal cells in j) and semi-quantitative analysis of t-STAT3 (positive epithelial cells in i) IHC stainings shown in g) (n≥6).

(f,h-j) Individual biological replicates are shown (f,h,j). Data are plotted as the means±SD and p-values were determined by unpaired two-tailed Student's t-tests (f,h), Mann-Whitney test (i) or ordinary one-way ANOVA with Tukey's multiple comparisons test (j).

**Supplementary Fig. 3**

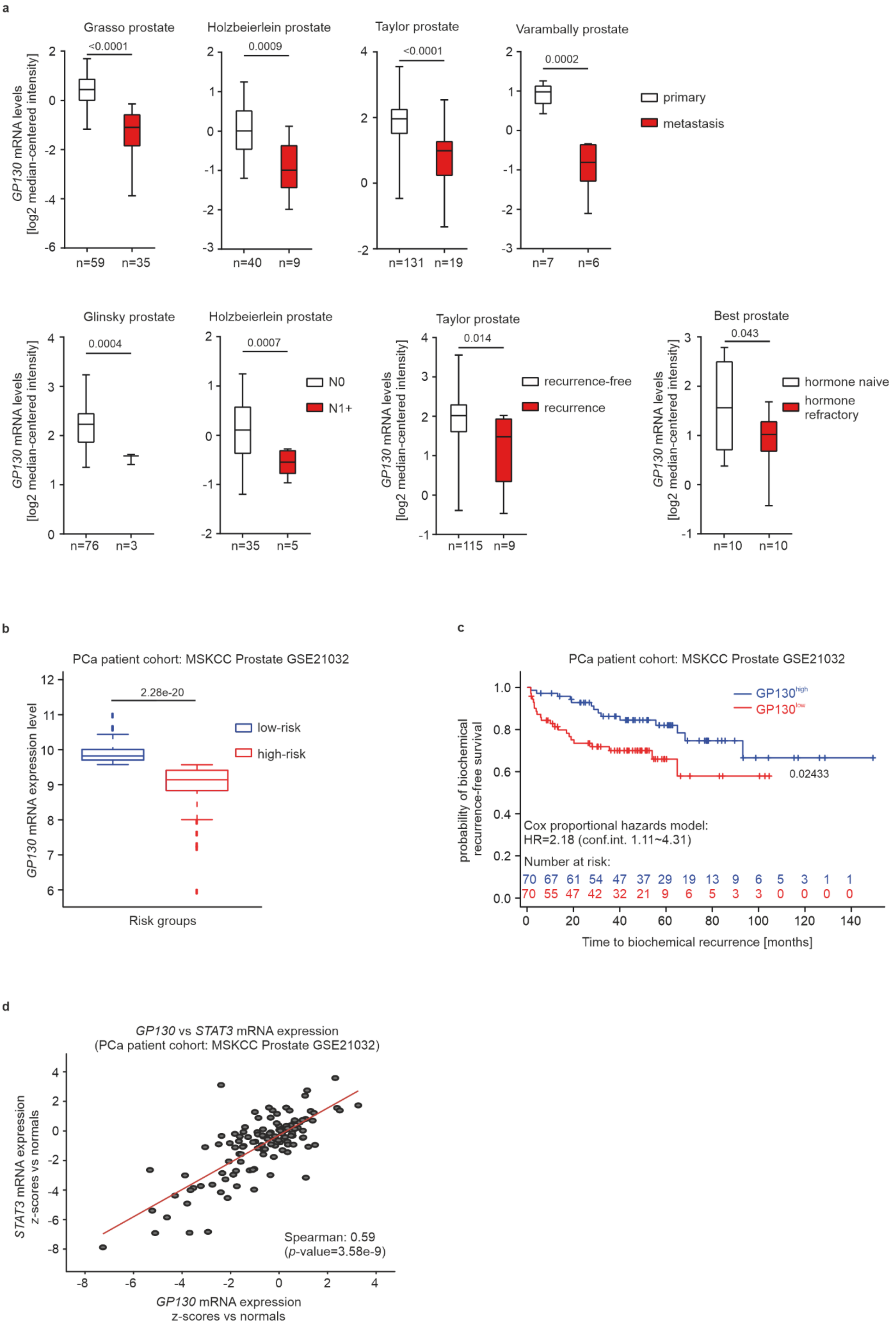

**Supplementary Fig. 3**

a) *GP130* mRNA expression levels of eight different data sets during PCa progression (recurrence, metastasis, advanced stage - metastatic spread to nearby lymph nodes (N category: N0 and N1+) and hormone refractory). Normalized data and statistical analyses were extracted from the Oncomine Platform. The respective prostate data set and n-numbers are indicated. Representation: boxes as interquartile range, horizontal line as the mean, whiskers as lower and upper limits.

b) Boxplots of *GP130* gene expression in risk groups of the MSKCC Prostate GSE21032 data set are shown. Risk groups were assessed by a median split of samples after Cox proportional hazards regression and ranking by their resulting prognostic index. P-values were determined by t-test. blue: low-risk group, red: high-risk group.

c) Kaplan-Meier plots showing time to biochemical recurrence in months for *GP130*$^{low}$ and *GP130*$^{high}$ risk groups (presented in b) of the MSKCC Prostate GSE21032 data set. Hazard ratio and p-value are estimated by a Cox-model using groups as covariate. blue: high *GP130* expressing, low-risk group, red: low *GP130* expressing, high-risk group. The blue and red numbers below horizontal axis represent the number of patients. Graphs (b-c) were generated by using the SurvExpress tool and the statistical analyses provided therein.

d) Spearman-correlation analysis of *GP130* and *STAT3* mRNA expression in MSKCC Prostate GSE21032 data set using cBioPortal analysis tool.

**Supplementary Fig. 4**

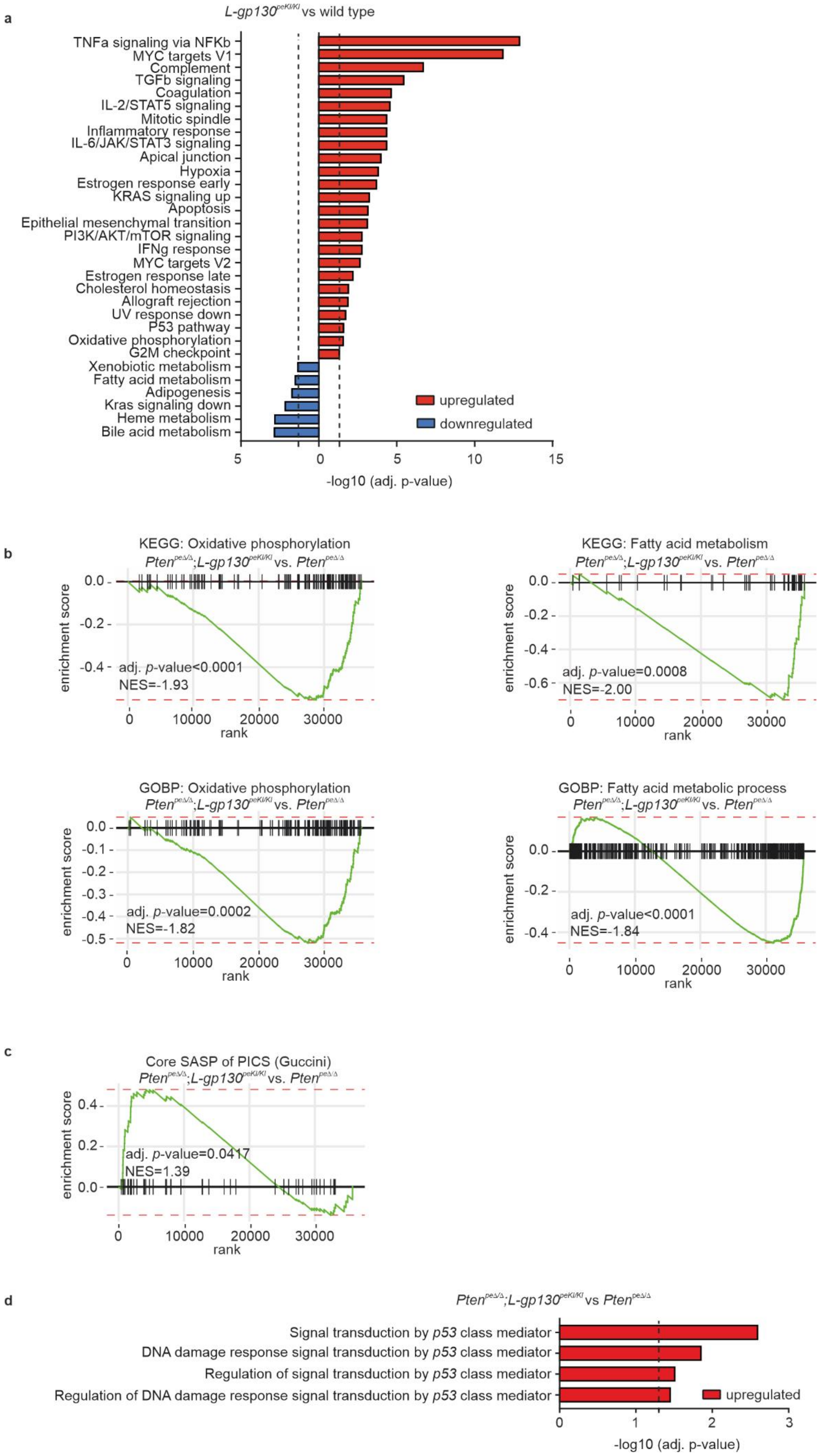

**Supplementary Fig. 4**

a) Fast pre-ranked gene set enrichment analysis (fGSEA) of significantly enriched HALLMARK gene sets with genes regulated in *L-gp130*$^{peKI/KI}$ compared to wild type prostate epithelial cells. Dotted line: adj. p-value (-log10(0.05)), blue: downregulated, red: upregulated;

b) Fast pre-ranked gene set enrichment analysis (fGSEA) of KEGG gene sets “KEGG: Oxidative phosphorylation” and “KEGG: Fatty acid metabolism” and Gene Ontology pathways, class Biological Processes (GOBP) gene sets “GOBP: Oxidative phosphorylation” and “GOBP: Fatty acid metabolism process” with genes regulated in *L-gp130*$^{peKI/KI}$ compared to wild type mice prostate epithelial cells. Genes sorted based on their Wald statistics are represented as vertical lines on the x-axis. NES: normalized enrichment score.

c) Fast pre-ranked gene set enrichment analysis (fGSEA) of the previously described core SASP gene signature upon PICS ”Core SASP of PICS (Guccini)“ with genes regulated in *Pten*$^{pe\Delta/\Delta}$;*L-gp130*$^{peKI/KI}$ compared to *Pten*$^{pe\Delta/\Delta}$ mice prostate epithelial cells. Genes sorted based on their Wald statistics are represented as vertical lines on the x-axis. NES: normalized enrichment score.

d) Fast pre-ranked gene set enrichment analysis (fGSEA) of selected Biological Processes from Gene Ontology pathways (GO-BP) gene sets related to p53 mediated signaling with genes significantly regulated in *Pten*$^{pe\Delta/\Delta}$;*L-gp130*$^{peKI/KI}$ compared to *Pten*$^{pe\Delta/\Delta}$ prostate epithelial cells. Dotted line: adj. p-value (-log10(0.05)), red: upregulated;

**Supplementary Fig. 5**

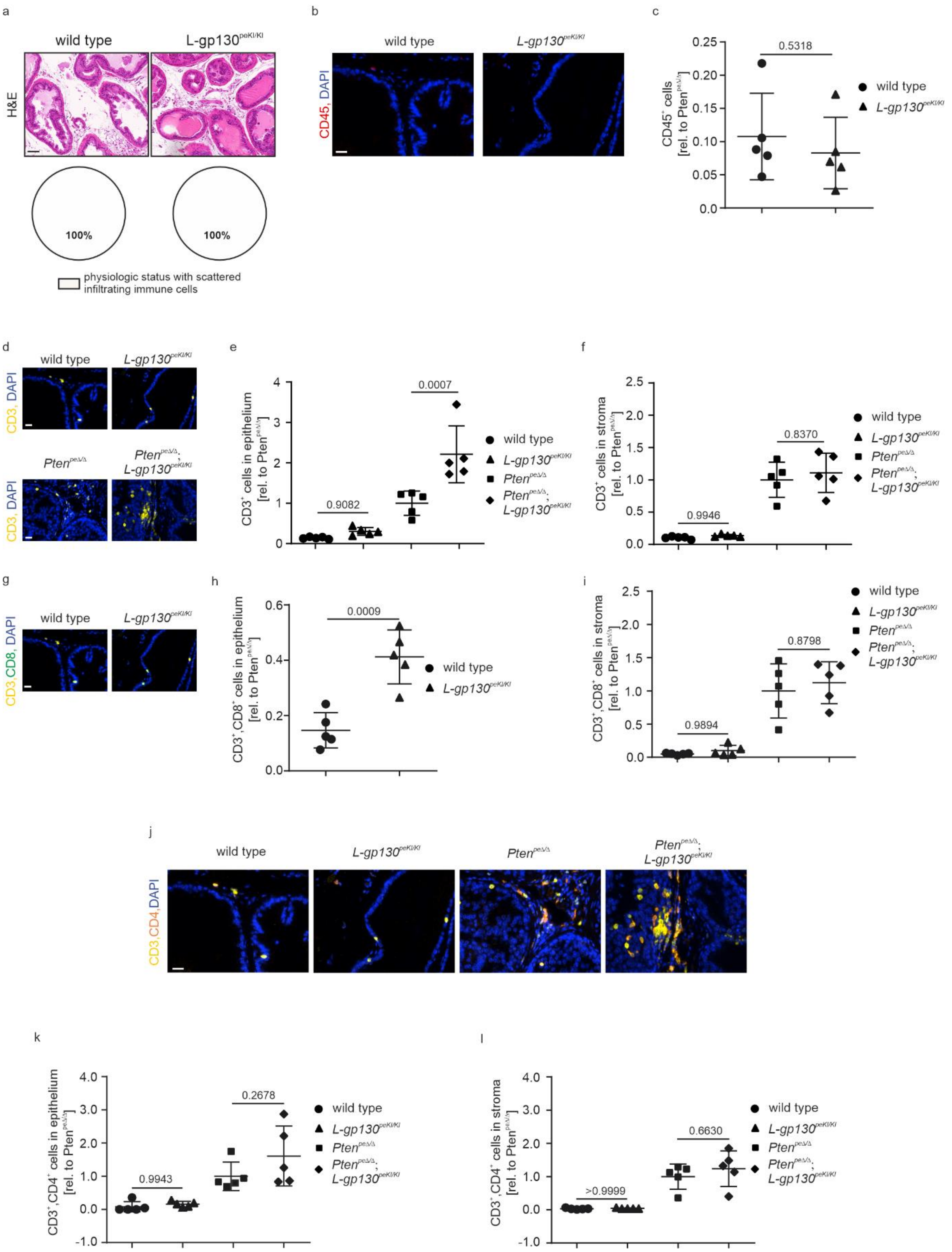

Supplementary Fig. 5 (continuation)

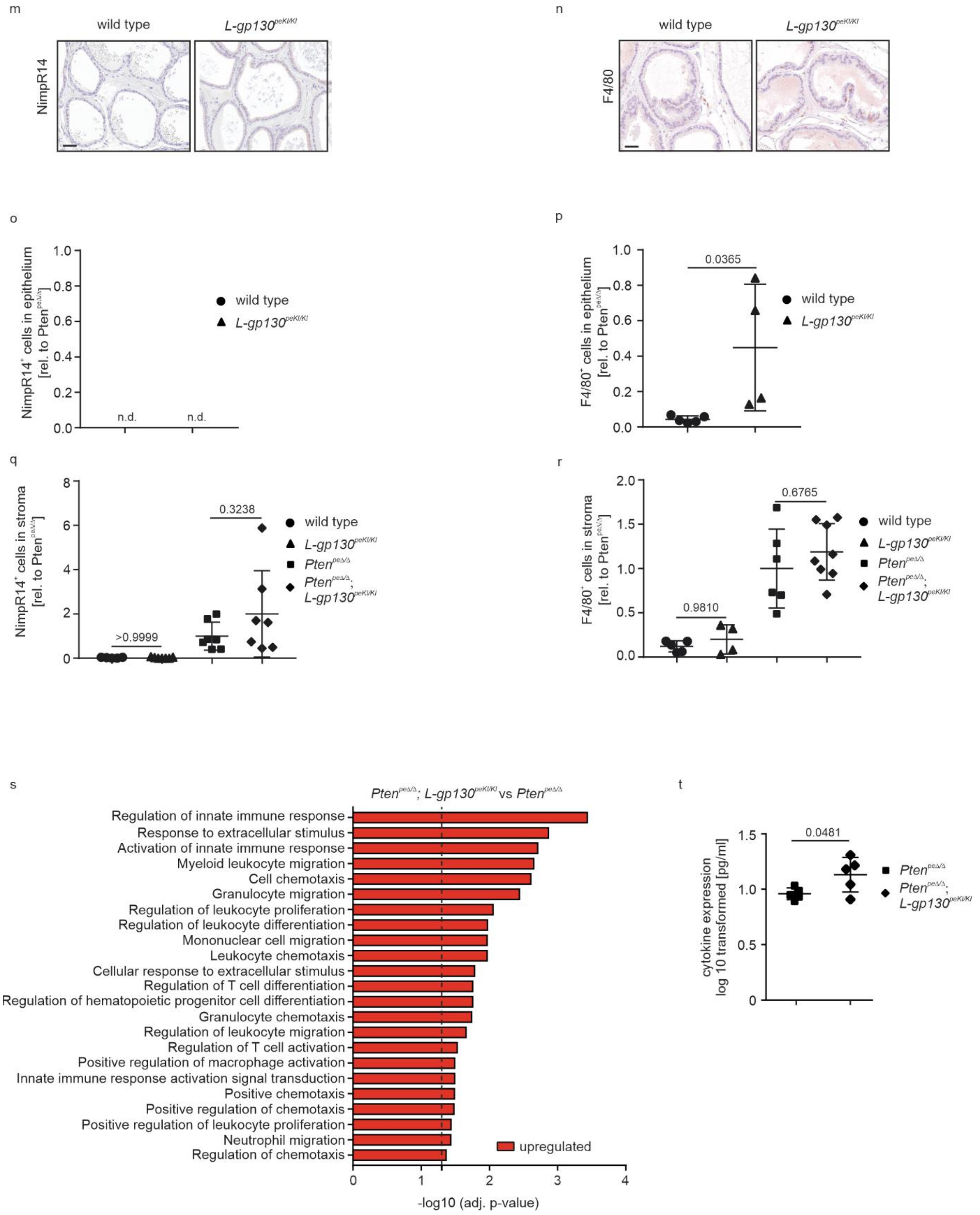


**Supplementary Fig. 5**

a) Representative pictures of H&E stains (upper panel) showing physiologic immune infiltrate and quantification of histopathological analysis (lower panel) of prostate tissue from wild type and *L-gp130*$^{peKI/KI}$ mice (n=9) in regards of infiltration (physiologic status with scattered infiltrating immune cell (white)). Scale bar: 60 µm.

b) Representative pictures of immunofluorescence (IF) staining of CD45 (red) and DAPI (blue) of mouse prostates with indicated genotypes. DAPI is used as a nuclear stain. Scale bar: 20 µm.

c) Quantification of $CD45^{+}$ cells of IF stainings shown in b) (n=5). A whole slide scan of stained prostate tissue per mouse was analyzed. The percentage of positive cells relative to *Pten*$^{pe\Delta/\Delta}$ was calculated.

d) Representative pictures of immunofluorescence (IF) staining of CD3 (yellow) and DAPI (blue) of mouse prostates with indicated genotypes. DAPI is used as a nuclear stain. Scale bar: 20 µm.

e-f) Quantification of $CD3^{+}$ cells in the prostate epithelium (e) and stroma (f) of IF stainings shown in d (n=5). A whole slide scan of stained prostate tissue per mouse was analyzed. The percentage of positive cells relative to *Pten*$^{pe\Delta/\Delta}$ was calculated.

g) Representative pictures of immunofluorescence (IF) staining of CD3 (yellow), CD8 (green) and DAPI (blue) of mouse prostates with indicated genotypes. DAPI is used as a nuclear stain. Scale bar: 20 µm.

h) Quantification of $CD3^{+}$;$CD8^{+}$ cells in the prostate epithelium of indicated genotypes (n=5) shown in g). A whole slide scan of stained prostate tissue per mouse was analyzed. The percentage of positive cells relative to *Pten*$^{pe\Delta/\Delta}$ was calculated.

i) Quantification of $CD3^{+}$;$CD8^{+}$ cells in the prostate stroma of IF stainings of indicated genotypes (n=5) shown in Fig. 5d and Supplementary Fig. 5g. A whole slide scan of stained prostate tissue per mouse was analyzed. The percentage of positive cells relative to *Pten*$^{pe\Delta/\Delta}$ was calculated.

j) Representative pictures of immunofluorescence (IF) staining of CD3 (yellow), CD4 (red) and DAPI (blue) of mouse prostates with indicated genotypes. DAPI is used as a nuclear stain. Scale bar: 20 µm.

k-l) Quantification of $CD3^{+}$;$CD4^{+}$ cells in the prostate epithelium (k) and stroma (l) of IF stainings shown in j (n=5). A whole slide scan of stained prostate tissue per mouse was analyzed. The percentage of positive cells relative to *Pten*$^{pe\Delta/\Delta}$ was calculated.

m-n) Representative pictures of immunofluorescence (IF) staining of NimpR14 (m) and F4/80 (n) of mouse prostates with indicated genotypes. Scale bar: 40 µm.

o-p) Quantification of $NimpR14^{+}$ (o) and $F4/80^{+}$ (p) cells in the prostate epithelium of IHC stainings shown in m-n (n≥4). A whole slide scan of stained prostate tissue per mouse was analyzed. The percentage of positive cells relative to *Pten*$^{pe\Delta/\Delta}$ was calculated. n.d.: not detected;

q-r) Quantification of $NimpR14^{+}$ (q) and $F4/80^{+}$ (r) cells in the prostate stroma of IHC stainings shown in Fig. 5f and Supplementary Fig. 5m-n (n≥4). A whole slide scan of stained prostate tissue per mouse was analyzed. The percentage of positive cells relative to *Pten*$^{pe\Delta/\Delta}$ was calculated.

s) Fast pre-ranked gene set enrichment analysis (fGSEA) of selected gene sets from Biological Processes from Gene Ontology pathways (GO-BP) related to the immune system with genes significantly regulated in *Pten*$^{pe\Delta/\Delta}$;*L-gp130*$^{peKI/KI}$ compared to *Pten*$^{pe\Delta/\Delta}$ prostate epithelial cells. Dotted line: adj. p-value (-log10(0.05)), red: upregulated;

t) Analysis of cytokine profile expression in the serum of *Pten*$^{pe\Delta/\Delta}$ and *Pten*$^{pe\Delta/\Delta}$;*L-gp130*$^{peKI/KI}$ mice (n=5).

(c,e-f,h-i,k-l,p-r,t) Individual biological replicates are shown. Data are plotted as the means±SD and p-values were determined by unpaired two-tailed Student's t-tests (c,h,p,t) or ordinary one-way ANOVA with Tukey's multiple comparisons test (e-f,i,k-l,q-r).

Supplementary Fig. 6

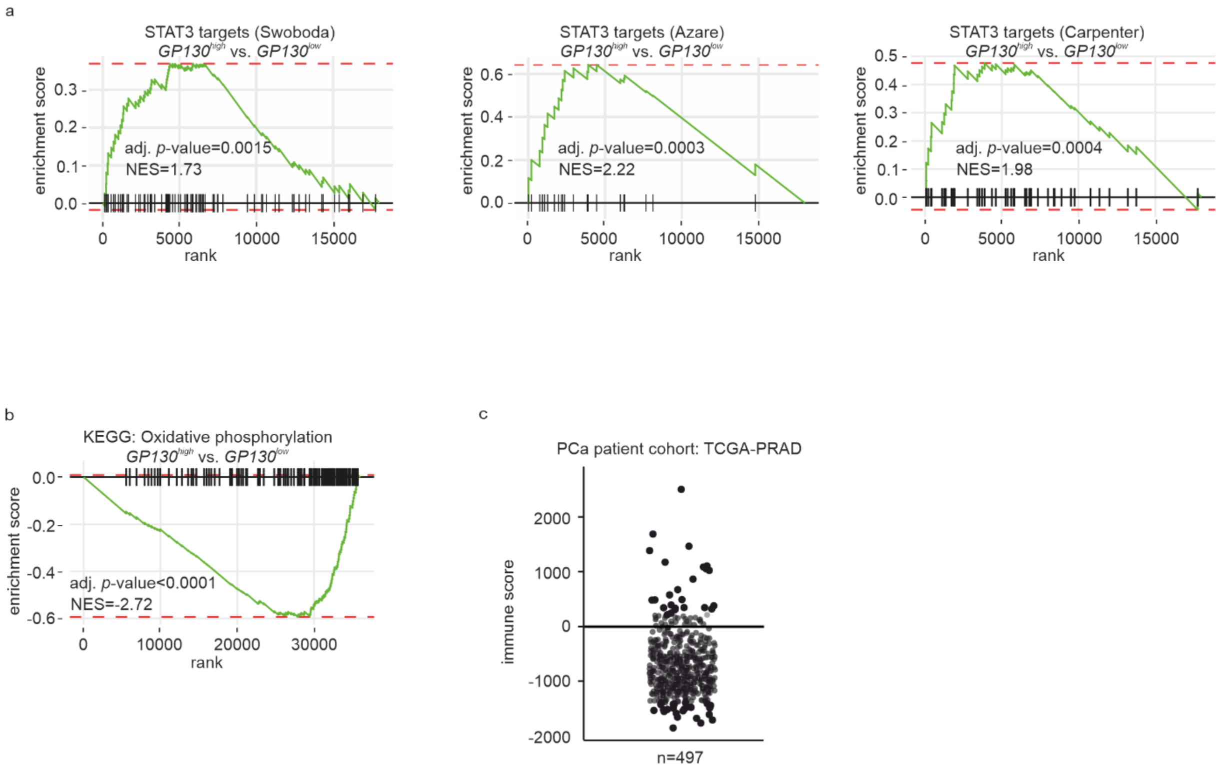


### Supplementary Fig. 6

a) Fast pre-ranked gene set enrichment analysis (fGSEA) of three previously published STAT3 target signatures (“STAT3 targets (Swoboda)”, “STAT3 targets (Azare)”, “STAT3 targets (Carpenter)”) with genes regulated in *GP130*$^{high}$ compared to *GP130*$^{low}$ expressing patients from the TCGA-PRAD data set. Genes sorted based on their Wald statistics are represented as vertical lines on the x-axis. NES: normalized enrichment score.

b) Fast pre-ranked gene set enrichment analysis (fGSEA) of the KEGG gene set “Oxidative phosphorylation” with genes regulated in *GP130*$^{high}$ compared to *GP130*$^{low}$ expressing patients from the TCGA-PRAD data set. Genes sorted based on their Wald statistics are represented as vertical lines on the x-axis. NES: normalized enrichment score.

c) Immune score from the ESTIMATE method for the TCGA-PRAD cohort. One dot represents one patient (n=497).

**Supplementary Table 1**

| Study of Origin | Sample ID | Mutation Type | Variant Type |
|---|---|---|---|
| Prostate Adenocarcinoma (TCGA, Firehose Legacy) | TCGA-YL-A8SB-01 | Missense_Mutation | SNP |
| Prostate Adenocarcinoma (TCGA, Firehose Legacy) | TCGA-ZG-A9LM-01 | Missense_Mutation | SNP |
| Prostate Adenocarcinoma (TCGA, Firehose Legacy) | TCGA-KC-A4BL-01 | Missense_Mutation | SNP |
| Prostate Adenocarcinoma (TCGA, Firehose Legacy) | TCGA-HC-A632-01 | Missense_Mutation | SNP |
| Prostate Adenocarcinoma (TCGA, Firehose Legacy) | TCGA-YL-A9WX-01 | Missense_Mutation | SNP |
| Prostate Adenocarcinoma (TCGA, Firehose Legacy) | TCGA-EJ-5521-01 | Missense_Mutation | SNP |
| Prostate Adenocarcinoma (TCGA, Firehose Legacy) | TCGA-HC-7230-01 | Missense_Mutation | SNP |
| Prostate Adenocarcinoma (TCGA, Firehose Legacy) | TCGA-HC-A631-01 | Missense_Mutation | SNP |
| Prostate Adenocarcinoma (TCGA, Firehose Legacy) | TCGA-KK-A8IG-01 | Missense_Mutation | SNP |
| Prostate Adenocarcinoma (TCGA, Firehose Legacy) | TCGA-YL-A8HK-01 | Missense_Mutation | SNP |
| Prostate Adenocarcinoma (TCGA, Firehose Legacy) | TCGA-KK-A6DY-01 | Missense_Mutation | SNP |
| Prostate Adenocarcinoma (TCGA, Firehose Legacy) | TCGA-EJ-5507-01 | Missense_Mutation | SNP |
| Prostate Adenocarcinoma (TCGA, Firehose Legacy) | TCGA-EJ-7315-01 | Missense_Mutation | SNP |
| Prostate Adenocarcinoma (TCGA, Firehose Legacy) | TCGA-VP-A872-01 | Missense_Mutation | SNP |
| Prostate Adenocarcinoma (TCGA, Firehose Legacy) | TCGA-EJ-5514-01 | Missense_Mutation | SNP |
| Prostate Adenocarcinoma (TCGA, Firehose Legacy) | TCGA-ZG-A9LB-01 | Missense_Mutation | SNP |
| Prostate Adenocarcinoma (TCGA, Firehose Legacy) | TCGA-G9-7510-01 | Missense_Mutation | SNP |
| Prostate Adenocarcinoma (TCGA, Firehose Legacy) | TCGA-V1-A8MJ-01 | Missense_Mutation | SNP |
| Prostate Adenocarcinoma (TCGA, Firehose Legacy) | TCGA-CH-5761-01 | Missense_Mutation | SNP |
| Prostate Adenocarcinoma (TCGA, Firehose Legacy) | TCGA-G9-7521-01 | Missense_Mutation | SNP |
| Prostate Adenocarcinoma (TCGA, Firehose Legacy) | TCGA-HI-7171-01 | Missense_Mutation | SNP |
| Prostate Adenocarcinoma (TCGA, Firehose Legacy) | TCGA-V1-A9O5-01 | Missense_Mutation | SNP |
| Prostate Adenocarcinoma (TCGA, Firehose Legacy) | TCGA-V1-A9O5-06 | Missense_Mutation | SNP |
| Prostate Adenocarcinoma (TCGA, Firehose Legacy) | TCGA-YL-A9WH-01 | Missense_Mutation | SNP |
| Prostate Adenocarcinoma (TCGA, Firehose Legacy) | TCGA-HC-8264-01 | Missense_Mutation | SNP |
| Prostate Adenocarcinoma (TCGA, Firehose Legacy) | TCGA-YL-A8HM-01 | Missense_Mutation | SNP |
| Prostate Adenocarcinoma (TCGA, Firehose Legacy) | TCGA-V1-A8WW-01 | Missense_Mutation | SNP |
| Prostate Adenocarcinoma (TCGA, Firehose Legacy) | TCGA-J4-A67L-01 | Missense_Mutation | SNP |
| Prostate Adenocarcinoma (TCGA, Firehose Legacy) | TCGA-EJ-A7NH-01 | Missense_Mutation | SNP |
| Prostate Adenocarcinoma (TCGA, Firehose Legacy) | TCGA-G9-A9S4-01 | Missense_Mutation | SNP |
| Prostate Adenocarcinoma (TCGA, Firehose Legacy) | TCGA-ZG-A9L1-01 | Missense_Mutation | SNP |
| Prostate Adenocarcinoma (TCGA, Firehose Legacy) | TCGA-HC-A9TH-01 | Missense_Mutation | SNP |
| Prostate Adenocarcinoma (TCGA, Firehose Legacy) | TCGA-EJ-8472-01 | Missense_Mutation | SNP |
| Prostate Adenocarcinoma (TCGA, Firehose Legacy) | TCGA-J9-A8CK-01 | Missense_Mutation | SNP |
| Prostate Adenocarcinoma (TCGA, Firehose Legacy) | TCGA-XJ-A9DI-01 | Missense_Mutation | SNP |
| Prostate Adenocarcinoma (TCGA, Firehose Legacy) | TCGA-HC-A48F-01 | Missense_Mutation | SNP |
| Prostate Adenocarcinoma (TCGA, Firehose Legacy) | TCGA-HC-8216-01 | Missense_Mutation | SNP |
| Prostate Adenocarcinoma (TCGA, Firehose Legacy) | TCGA-G9-6499-01 | Missense_Mutation | SNP |
| Prostate Adenocarcinoma (TCGA, Firehose Legacy) | TCGA-YL-A9WK-01 | Missense_Mutation | SNP |
| Prostate Adenocarcinoma (TCGA, Firehose Legacy) | TCGA-VP-A876-01 | Missense_Mutation | SNP |
| Prostate Adenocarcinoma (TCGA, Firehose Legacy) | TCGA-G9-A9S0-01 | Missense_Mutation | SNP |
| Prostate Adenocarcinoma (TCGA, Firehose Legacy) | TCGA-HC-7213-01 | Splice_Site | SNP |
| Prostate Adenocarcinoma (TCGA, Firehose Legacy) | TCGA-KC-A7FA-01 | Splice_Site | SNP |
| Prostate Adenocarcinoma (TCGA, Firehose Legacy) | TCGA-KK-A7AU-01 | Splice_Site | SNP |
| Prostate Adenocarcinoma (TCGA, Firehose Legacy) | TCGA-2A-A8VT-01 | Splice_Site | SNP |
| Prostate Adenocarcinoma (TCGA, Firehose Legacy) | TCGA-J9-A52B-01 | Splice_Site | DEL |
| Prostate Adenocarcinoma (TCGA, Firehose Legacy) | TCGA-KK-A8II-01 | Splice_Site | DEL |

| | | | |
|---|---|---|---|
| Prostate Adenocarcinoma (TCGA, Firehose Legacy) | TCGA-YL-A9WY-01 | Missense_Mutation | SNP |
| Prostate Adenocarcinoma (TCGA, Firehose Legacy) | TCGA-ZG-A9L0-01 | Nonsense_Mutation | SNP |
| Prostate Adenocarcinoma (TCGA, Firehose Legacy) | TCGA-KK-A7B4-01 | Frame_Shift_Ins | INS |
| Prostate Adenocarcinoma (TCGA, Firehose Legacy) | TCGA-ZG-A9M4-01 | Nonsense_Mutation | SNP |
| Prostate Adenocarcinoma (TCGA, Firehose Legacy) | TCGA-HC-7742-01 | Frame_Shift_Del | DEL |
| Prostate Adenocarcinoma (TCGA, Firehose Legacy) | TCGA-V1-A9ZI-01 | Frame_Shift_Del | DEL |
| Prostate Adenocarcinoma (TCGA, Firehose Legacy) | TCGA-EJ-8472-01 | Missense_Mutation | SNP |
| Prostate Adenocarcinoma (TCGA, Firehose Legacy) | TCGA-VP-A87D-01 | Frame_Shift_Del | DEL |
| Prostate Adenocarcinoma (TCGA, Firehose Legacy) | TCGA-HC-A9TE-01 | Frame_Shift_Del | DEL |
| Prostate Adenocarcinoma (TCGA, Firehose Legacy) | TCGA-EJ-7781-01 | Frame_Shift_Del | DEL |
| Prostate Adenocarcinoma (TCGA, Firehose Legacy) | TCGA-YL-A8HL-01 | Missense_Mutation | SNP |
| Prostate Adenocarcinoma (TCGA, Firehose Legacy) | TCGA-YL-A8SJ-01 | Frame_Shift_Ins | INS |
| Prostate Adenocarcinoma (TCGA, Firehose Legacy) | TCGA-EJ-5525-01 | Frame_Shift_Del | DEL |
| Prostate Adenocarcinoma (TCGA, Firehose Legacy) | TCGA-ZG-A9M4-01 | Frame_Shift_Del | DEL |
| Prostate Adenocarcinoma (TCGA, Firehose Legacy) | TCGA-KC-A7F6-01 | Frame_Shift_Del | DEL |
| Prostate Adenocarcinoma (TCGA, Firehose Legacy) | TCGA-XK-AAIW-01 | Missense_Mutation | SNP |
| Prostate Adenocarcinoma (TCGA, Firehose Legacy) | TCGA-XK-AAIW-01 | Missense_Mutation | SNP |

### Supplementary Table 2

**qRT-qPCR primers**

| Primer Name | Primer Sequence 5'-3' |
|---|---|
| mRT cyclophilinA fw | TCGAGCTCTGAGCACTGGAG |
| mRT cyclophilinA rv | CATTATGGCGTGTAAAGTCACCA |
| mRT Hprt fw | GCTGGTGAAAAGGACCTCT |
| mRT Hprt rv | CACAGGACTAGAACACCTGC |
| 18S fw | GCCCGAAGCGTTTACTTTGA |
| 18S rv | TCCATTATTCCTAGCTGCGGTATC |
| mRT p19ARF fw | TGGTCACTGTGAGGATTCAGC |
| mRT p19ARF rv | CGTGAACGTTGCCCATCATC |
| mRT Gp130 fw | TGGCCTAATGTTCCTGATCC |
| mRT Gp130 rv | GCCGTCCGAGTACATTTGAT |

**Genotyping primers**

| Primer Name | PrimerSequence 5'-3' |
|---|---|
| Cre fw | CGGTCGATGCAACGAGTGATGAGG |
| Cre rv | CCAGAGACGGAAATCCATCGCTCG |
| Pten fw | CTCCTCTACTCCATTCTTCCC |
| Pten rv | ACTCCCACCAATGAACAAAC |
| Pten KO fw | GTCACCAGGATGCTTCTGAC |
| Pten KO rv | ACTATTGAACAGAATCAACCC |
| L-gp130 wild type fw | TGTCGCAAATTAACTGTGAATC |
| L-gp130 transgene fw | GATATGAAGTACTGGGCTCTT |
| L-gp130 wild type+transgene rv | AAAGTCGCTCTGAGTTGTTATC |
| L-gp130 KI fw | TGTCATACTTATCCTGTCCCTTTTT |
| L-gp130 KI rv | CAGAAGAGTTGTCAATAGGAATGCT |

**Supplementary Table 3**

**Genotyping protocol**

| Step | Temp. [°C] | Time | No. of cycles |
|---|---|---|---|
| Initial Denaturation | 94 | 2 min | 1 |
| Denaturation | 94 | 20 sec | |
| Annealing | 55 | 30 sec | 32 |
| Extension | 72 | 2 min | |
| Final Extension | 72 | 10 min | 1 |
| Storage | 12 | for ever | |

### Supplementary Table 4

| Primary Antibody, Company, Product number | Application | Pretreatment | Autostainer/manually | Dilution | Seondary Antibody |
|---|---|---|---|---|---|
| **phospho-AKT**<br>Cell Signaling<br>4060 | IHC | citrate buffer pH6 | Lab Vision 360 Thermo Scientific | 1:200 | secondary antibody conjugated to enzyme labelled polymer (Bright Vision, Rabbit HRP KL DPVR 110 HRP) |
| **EpCAM**<br>Elab Sci<br>E-AB-70132 | IHC | EDTA buffer pH8 | Lab Vision 360 Thermo Scientific | 1:1300 | secondary antibody conjugated to enzyme labelled polymer (Bright Vision, Rabbit HRP KL DPVR 110 HRP) |
| **phosphoY-STAT3**<br>Cell Signaling<br>9145 | IHC | Tris-EDTA pH9 | Lab Vision 360 Thermo Scientific | 1:100 | secondary antibody conjugated to enzyme labelled polymer (Bright Vision, Rabbit HRP KL DPVR 110 HRP) |
| **total-STAT3**<br>Cell Signaling<br>12640 | IHC | Tris-EDTA pH9 | Lab Vision 360 Thermo Scientific | 1:3000 | secondary antibody conjugated to enzyme labelled polymer (Bright Vision, Rabbit HRP KL DPVR 110 HRP) |
| **Ki67**<br>Cell Signaling<br>12202 | IHC | citrate buffer pH6 | Lab Vision 360 Thermo Scientific | 1:1000 | secondary antibody conjugated to enzyme labelled polymer (Bright Vision, Rabbit HRP KL DPVR 110 HRP) |
| **PML**<br>Millipore<br>MAB3738 | IHC | Citrate buffer, pH6, steamer | manually | 1:50 | UltraTek Anti-Polyvalent (ScyTek laboratories, AFN600) |
| **F4/80**<br>Cell Signaling<br>70076 | IHC | citrate buffer pH6 | Lab Vision 360 Thermo Scientific | 1:1000 | Biotinylated anti Rabbit (Vector, BA-1000) |
| **NimpR14**<br>Abcam<br>Ab2557 | IHC | Pronase | Lab Vision 360 Thermo Scientific | 1:1000 | Biotinylated anti Rat (Vector, BA-4001) |
| **CD3**<br>DAKO<br>A0452 | IHC | citrate buffer pH6 | Lab Vision 360 Thermo Scientific | 1:1000 | secondary antibody conjugated to enzyme labelled polymer (Bright Vision, Rabbit HRP KL DPVR 110 HRP) |

| | | | | | |
|---|---|---|---|---|---|
| **CD45**<br>Abcam plc<br>ab208022 | IF | Bake: 72°C 30min., Dewax with dewax solution (Leica, AR9222) | Leica, BondRX | 1:7000 | EnVision, HRP Labelled AB - Anti Rabbit (DAKO, K400311-2) |
| **CD3**<br>Abcam plc<br>ab135372 | IF | Bake: 72°C 30min., Dewax with dewax solution (Leica, AR9222) | Leica, BondRX | 1:100 | EnVision, HRP Labelled AB - Anti Rabbit (DAKO, K400311-2) |
| **CD8 alpha**<br>Cell Signaling<br>98941S | IF | Bake: 72°C 30min., Dewax with dewax solution (Leica, AR9222) | Leica, BondRX | 1:500 | EnVision, HRP Labelled AB - Anti Rabbit (DAKO, K400311-2) |
| **CD4**<br>Abcam plc<br>ab183685 | IF | Bake: 72°C 30min., Dewax with dewax solution (Leica, AR9222) | Leica, BondRX | 1:300 | EnVision, HRP Labelled AB - Anti Rabbit (DAKO, K400311-2) |
| **ZSGreen**<br>Clontech<br>632474 | IF | - | Lab Vision 360 Thermo Scientific | 1:250 | Rabbit Alexa Fluor 594 (Invitrogen, A11037) |
| **phospho-AKT**<br>Cell Signaling<br>4060 | WB | - | - | according to manufacturer’s instructions | Anti rabbit IgG, HRP linked (Cell Signaling, 7074) |
| **total-AKT**<br>Cell Signaling<br>4691 | WB | - | - | according to manufacturer’s instructions | Anti rabbit IgG, HRP linked (Cell Signaling, 7074) |
| **phosphoY-STAT3**<br>Cell Signaling<br>9145 | WB | - | - | according to manufacturer’s instructions | Anti rabbit IgG, HRP linked (Cell Signaling, 7074) |
| **total-STAT3**<br>Cell Signaling<br>9139 | WB | - | - | according to manufacturer’s instructions | Anti mouse IgG, HRP-linked AB (GE Healthcare, Amersham, NXA931) |
| **p53**<br>Cell Signaling<br>2524 | WB | - | - | according to manufacturer’s instructions | Anti mouse IgG, HRP-linked AB (GE Healthcare, Amersham, NXA931) |
| **β-ACT**<br>Cell Signaling<br>3700/4967 | WB | - | - | according to manufacturer’s instructions | Anti mouse IgG, HRP-linked AB (GE Healthcare, Amersham, NXA931)/Anti rabbit IgG, HRP linked (Cell Signaling, 7074) |
